# TBC1D23-AAVR Interaction Drives Endosome-to-TGN Trafficking Required for rAAV Transduction

**DOI:** 10.64898/2026.06.02.729721

**Authors:** Xiujuan Zhang, Ariful Habib, Shane McFarlin, Donovan Richart, Soo Yeun Park, Fang Cheng, Jinxi Wang, Ziying Yan, Jianming Qiu

**Affiliations:** Department of Microbiology, Molecular Genetics and Immunology University of Kansas Medical Center, Kansas City, KS 66160, USA; Division of Pulmonary, Allergy and Critical Care Medicine Department of Medicine, University of Alabama at Birmingham Birmingham, AL 35294, USA; Department of Orthopedic Surgery and Sports Medicine University of Kansas Medical Center, Kansas City, KS 66160, USA

**Keywords:** AAV, AAVR, TBC1D23, retrograde transport, transduction

## Abstract

The adeno-associated virus receptor (AAVR; also known as KIAA0319L) is the proteinaceous receptor required for the transduction of multi-serotype adeno-associated viruses (AAVs). While the extracellular polycystic kidney disease (PKD) domains of AAVR directly bind AAV capsids, the function of its C-terminal cytosolic domain (AAVR-C) in governing AAV internalization and intracellular trafficking remains undefined. Using targeted pulldown of AAVR-interacting host proteins with a glutathione S-Transferase (GST)-fused AAVR-C recombinant protein (GST-AAVR-C) as a bait, we identified that TBC1 domain family member 23 (TBC1D23), a specialized intracellular trafficking adaptor and bridging factor, binds AAVR via the AAVR-C and mediates AAV’s endosome-to-*trans*-Golgi network (TGN) transport. Biolayer interferometry (BLI) demonstrates nanomolar-affinity binding between AAVR-C and the C-terminal scaffold domain of TBC1D23 (TBC1D23-C). CRISPR-mediated knockout of *TBC1D23* severely impairs rAAV transduction across multiple human cell types, including polarized human airway epithelium (HAE). Loss of TBC1D23 disrupts convergence of internalized AAV capsids at the TGN, leading to diminished nuclear import of AAV vectors. Mutational analyses of AAVR-C reveal that its acidic residue cluster is required for TBC1D23 binding and AAV retrograde transport from the endosome to the TGN. Together, these findings define TBC1D23 as a receptor-proximal trafficking module that couples AAV-AAVR engagement to vesicle transport, revealing a missing core regulatory step in AAV retrograde transport that is essential for productive rAAV transduction.

**Significance:** Recombinant adeno-associated viruses (rAAVs) are widely used gene delivery vectors, yet the intracellular trafficking steps that enable productive transduction remain incompletely defined. We identify TBC1D23 as a critical host adaptor that directly binds the cytosolic tail of the AAV receptor (AAVR) and mediates retrograde transport of AAV vectors from endosomes to the *trans*-Golgi network (TGN). Disruption of the AAVR acidic residue cluster abolishes TBC1D23 binding, blocks TGN trafficking, impairs nuclear import, and severely reduces transduction across multiple serotypes and in polarized human airway epithelium. These findings reveal a receptor-proximal trafficking checkpoint essential for rAAV gene delivery and define a host–vector interface with potential for rational vector engineering and therapeutic modulation.

## Introduction

Adeno-associated viruses (AAVs) are small, non-enveloped parvoviruses widely used gene delivery vectors. Their favorable safety profile, ability to transduce both dividing and non-dividing cells, and the capacity for sustained transgene expression have established recombinant AAV (rAAV) vectors as a leading platform for human gene therapy (1). Despite these advantages, clinical applications are often constrained by the need for high-dose administration, which increases the risk of host immune response against the viral capsid and genome, even though AAVs are generally considered low in immunogenicity (2,3). Although rAAVs are widely used in clinical settings with several licensed gene therapies approved for the treatment of rare monogenic disorders (1), the mechanisms governing AAV entry, intracellular trafficking, nuclear import, and double-stranded (ds)DNA genome synthesis and recombination remain incompletely understood (4,5). A more detailed mechanistic understanding of these processes could inform strategies to improve transduction efficiency and safety via either modulating the vector-host interactions or rationally engineering vector tropism.

AAV entry proceeds through a two-step receptor engagement process: beginning with attachment to cell-surface glycans and followed by binding to a secondary proteinaceous receptor that mediates endocytosis and subsequent intracellular trafficking to the perinuclear region (6,7). KIAA0319L was identified as a broadly acting proteinaceous receptor for multi-serotype AAVs and was subsequently named the AAV receptor (AAVR) (8). AAVR is a type I transmembrane protein comprising an ectodomain composed of an N-terminal signal peptide (SP), an MANEC (Motif at N Terminus with Eight Cysteines) domain, five extracellular polycystic kidney disease (PKD) domains (PKD1-5) and a single transmembrane (TM) region, followed by a short cytoplasmic C-terminal tail (AAVR-C) containing motifs implicated in endosomal sorting (6,8) (**Fig. S2A**). While PKD1-5 directly bind AAV capsids, notably PKD1 for AAV5 and PKD2 for AAV2 (6,9), deletion of AAVR-C results in increased cell surface expression but impaired internalization and an inability to rescue rAAV transduction in AAVR knockout (KO) cells (8). These facts position AAVR as a central determinant of AAV endocytosis and intracellular trafficking post-attachment. However, how the AAVR-C interacts with host proteins and engages intracellular trafficking machinery to mediate these processes remains unclear.

Although transiently expressed at the plasma membrane, AAVR predominantly localizes to the *trans*-Golgi network (TGN) and cycles between the cell surface and the TGN (8). The prevailing model proposes that AAVR functions as a sorting receptor that captures AAV capsids at the plasma membrane, mediates their endocytosis into endosomal compartments, and directs them to the retrograde transport pathway toward the TGN. This route is thought to be critical for capsid processing (10), including exposure of the VP1 unique region (VP1u), which harbors a phospholipase A₂ domain essential for membrane vesicle escape, later during trafficking to perinuclear regions (11). Despite the central role of AAVR in this pathway, the host machinery that coordinates AAVR’s endocytosis and intracellular trafficking between the cell surface, endosomes, and the TGN remains undefined. This machinery likely includes adaptors and vesicle-tethering factors that link AAVR to the retrograde transport pathway (7).

To identify host factors that mediate AAVR-dependent AAV intracellular trafficking, we carried out a glutathione S-transferase (GST) pulldown assay using GST-fused AAVR-C to isolate interacting proteins, followed by quantitative mass spectrometry (qMS). This approach identified that TBC1D23 (TBC1 domain family member 23) specifically interacts with AAVR-C, and that this interaction is critical for AAVR-mediated trafficking of AAV toward productive transduction. TBC1D23 is a Golgi-localized, catalytically inactive TBC-domain protein that functions as a vesicle-tethering adaptor mediating endosome-to-TGN trafficking. By bridging endosome-associated WASH/FAM21 complexes to TGN-localized golgins (golgin-97 and-245), TBC1D23 enables the retrograde transport of selective cargoes (12–14). At the TGN, TBC1D23 further cooperates with the WDR11–FAM91A1 complex, a TGN-localized vesicle capture and stabilization module that promotes efficient docking and retention of incoming endosome-derived carriers prior to membrane fusion (13–15). TBC1D23 scaffolds Golgi-specific signaling modules, including carboxypeptidase D (CPD), TANK-binding kinase 1 (TBK1), and liver kinase B1 (LKB1), highlighting its broader role in coordinating spatiotemporally restricted trafficking and signaling events at the TGN (14–17). Disruption of TBC1D23, therefore, impairs both retrograde vesicle transport and Golgi-associated signaling.

In this study, we demonstrate that the C-terminal tail of AAVR directly binds the C-terminal scaffold domain of TBC1D23 with nanomolar affinity, and that this interaction is essential for AAV intracellular trafficking and transduction. Using CRISPR-mediated gene KO, AAVR complementation, confocal imaging, and endosome immunoprecipitation (endo-IP), we confirmed TBC1D23 as a receptor-proximal trafficking factor that couples AAV–AAVR engagement to endosome-to-TGN transport.

## Results

### AAVR is required for efficient AAV internalization

Although AAVR has been identified as an essential receptor for multi-serotype AAVs, the fundamental questions regarding whether AAVR interacts with AAV at the plasma membrane and mediates AAV internalization remain elusive. We analyzed the colocalization of AAV5 capsids with AAVR during the early stages of transduction in HEK293 cells. We found AAV5 capsids colocalized with AAVR on the plasma membrane of wild-type (WT) cells stained with Maackia amurensis lectin (MAL) II at 2 h post-transduction (hpt); in contrast, little or no AAV5 capsids were detected in AAVR-KO HEK293 cells (**Fig. 1A**). We further quantified vector genomes that were internalized into the cells at 2 hpt using qPCR. We found that the internalized vector genomes were significantly reduced by 98.0% in AAVR-KO cells (**Fig. 1B**). Moreover, at 4 hpt, internalized rAAV5 particles showed partial colocalization with AAVR and syntaxin 5 (STX5)-positive endosomal compartments in WT cells (**Fig. S1A**). At 6 hpt, rAAV5 capsids accumulated in the perinuclear region and partially colocalized with AAVR and TGN46- positive compartments in WT cells (**Fig. S1B**). No capsid signals or AAVR were detected in AAVR-KO cells Taken together, these results strongly support the conclusion that AAVR is required for efficient AAV internalization and may also mediate post-entry intracellular trafficking.

**Fig. 1.**
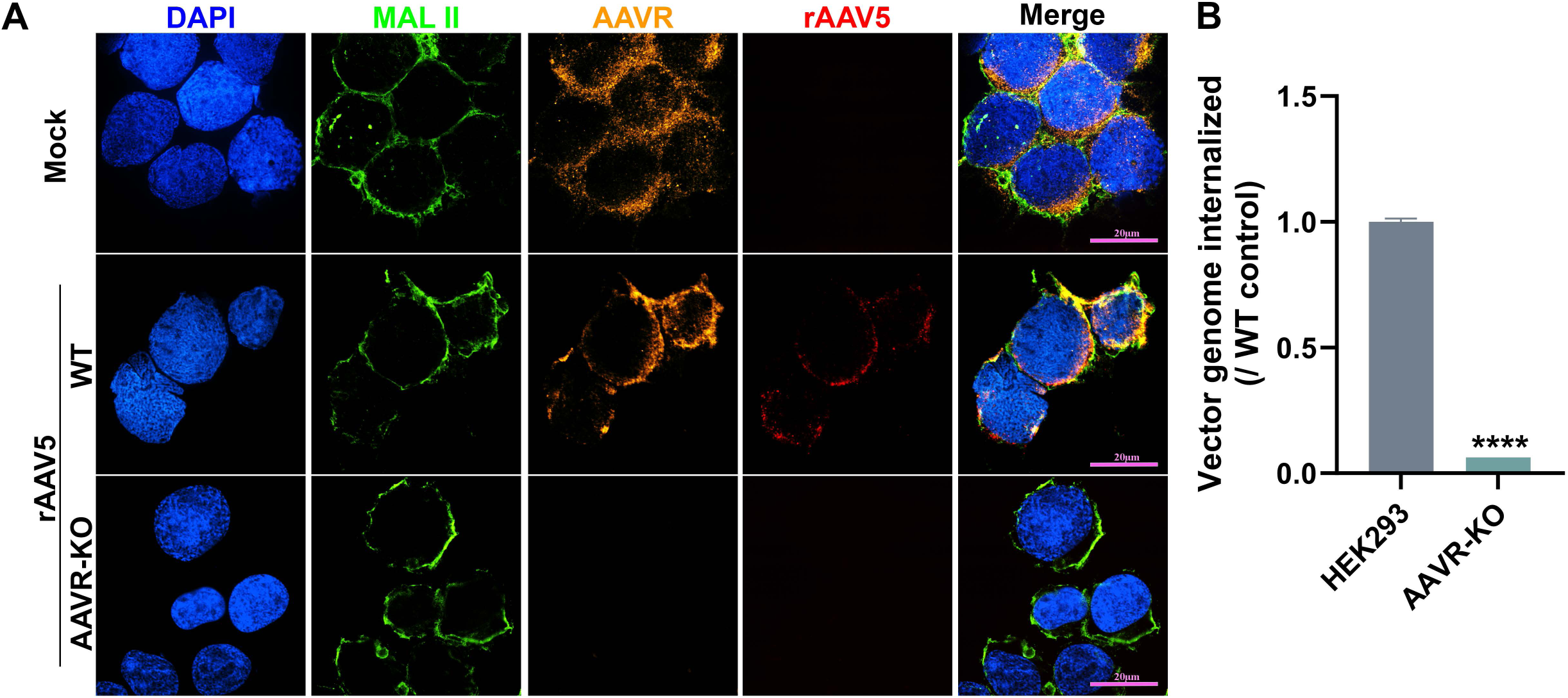
AAVR colocalizes with cell-surface sialic acid and TGN/Golgi-associated vesicles during rAAV5 entry. Wild-type (WT) or AAVR-knockout (AAVR-KO) HEK293 cells were incubated with rAAV5 at an MOI of 50K DRP/cell at 4°C for 1 h, followed by incubation at 37°C for 2 h. Cells were fixed and stained as indicated. Nuclei were counterstained with DAPI (blue). rAAV5 capsids are shown in red and AAVR in orange. **(A) AAVR localizes on plasma membrane with AAV capsid.** At 2 h post-transduction (hpt), cell-surface α2,3-linked sialic acid was detected using biotinylated MAL II lectin (green) prior to permeabilization. **(B)** Quantification of internalized vector genomes in WT and AAVR-KO HEK293 cells at 2 hpt. Data are presented as mean ± SD. ****P < 0.0001.

### TBC1D23 is identified as a cytosolic trafficking adaptor that binds the AAVR C-terminal tail

To explore the role of AAVR in AAV intracellular trafficking, we sought to identify host proteins that interact with the cytosolic tail of the AAVR (AAVR-C). We performed affinity purification using GST-fused AAVR-C (GST-AAVR-C) (**Fig. S2A&B**), followed by quantitative mass spectrometry (qMS). GST-AAVR-C-coated glutathione resin was incubated with lysates of HEK293 cells, and the proteins retained on the GST–AAVR-C resin or GST alone-coated resin were eluted and analyzed by liquid chromatography-tandem mass spectrometry (LC-MS/MS) (**Table S1**). Comparative proteomic analysis revealed a subset of proteins significantly enriched in the GST–AAVR-C pulldown relative to the GST control (**Fig. S2C**). Gene ontology (GO) analysis of the most highly enriched candidates revealed enrichment of proteins involved in intracellular membrane trafficking and vesicular transport, with TBC1D23, COPG1, and ASAP1 among the most prominent candidates (**Fig. 2A**).

**Fig. 2.**
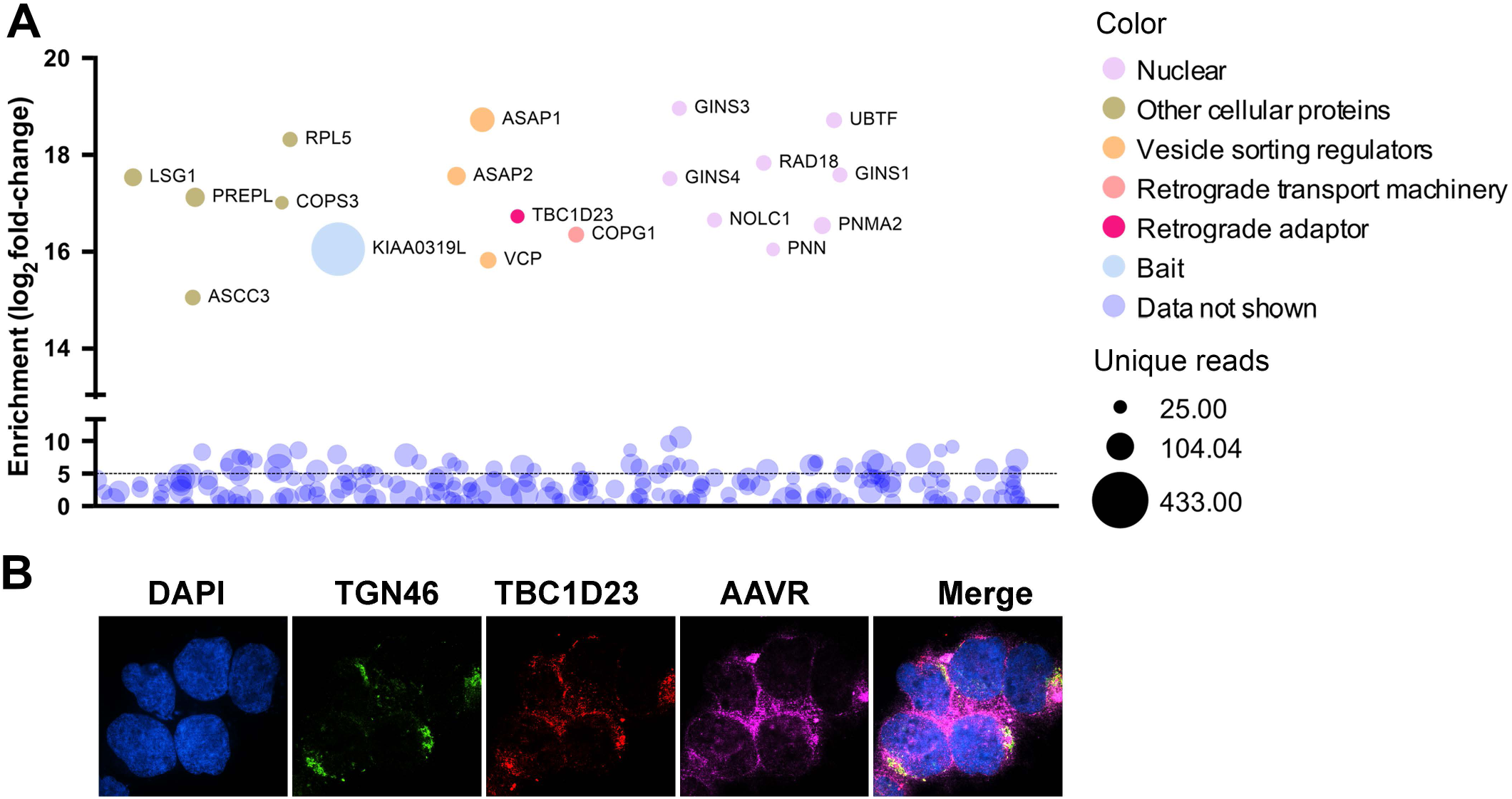
Identification of TBC1D23 as a cytosolic interactor of KIAA0319L (AAVR). (A) Gene Ontology (GO) analysis. The proteins enriched in GST-AAVR-C pull-downs (with ≥25 unique peptides, fold-change >2^10^, and *p* < 0.05) were considered significantly enriched) were grouped by subcellular localization and biological function. Proteins associated with the Golgi apparatus and membrane trafficking pathways are prominently represented. Bubble size reflects the number of unique peptides detected. TBC1D23 and COPG1 are highlighted among the significantly enriched Golgi-associated factors. **(B) TBC1D23 association with the AAVR.** HEK293 cells were stained for AAVR, TBC1D23, and the TGN marker TGN46, with nuclei labeled by DAPI. Images were taken under a confocal microscope (CSU-W1 SoRa, Nikon) with a 60× objective. Merged images indicate overlapping localization of AAVR with TBC1D23 within TGN46-associated compartments.

To validate the candidates identified by qMS, we repeated the GST pulldown and analyzed the precipitated proteins by immunoblotting. The results confirmed that TBC1D23, COPG1, ASAP1 and COPS3 were specifically enriched in the GST–AAVR-C pulldown but not in the GST control (**Fig. S3A&B**). We next use CRISPR-Cas9 to generate TBC1D23 knockout (KO) cells (**Fig. S4A**) and utilized siRNA to knock down COPG1, ASAP1 and COPS3 (**Fig. S3C**). We next evaluated AAV transduction in these gene KO or knockdown (KD) cells. TBC1D23-KO markedly impaired rAAV2.5T transduction and led to ∼95% reduction in mCherry-positive cells and a 73% decrease in firefly luciferase (fLuc) activity (**Fig. 3A&B**), whereas KD of COPG1, ASAP1, or COPS3 had little or no effect (**Fig. S3D**). Confocal microscopy further demonstrated that endogenous TBC1D23 colocalizes with AAVR in TGN46-positive compartments (**Fig. 2B**), supporting its physiological interaction at the TGN.

**Fig. 3.**
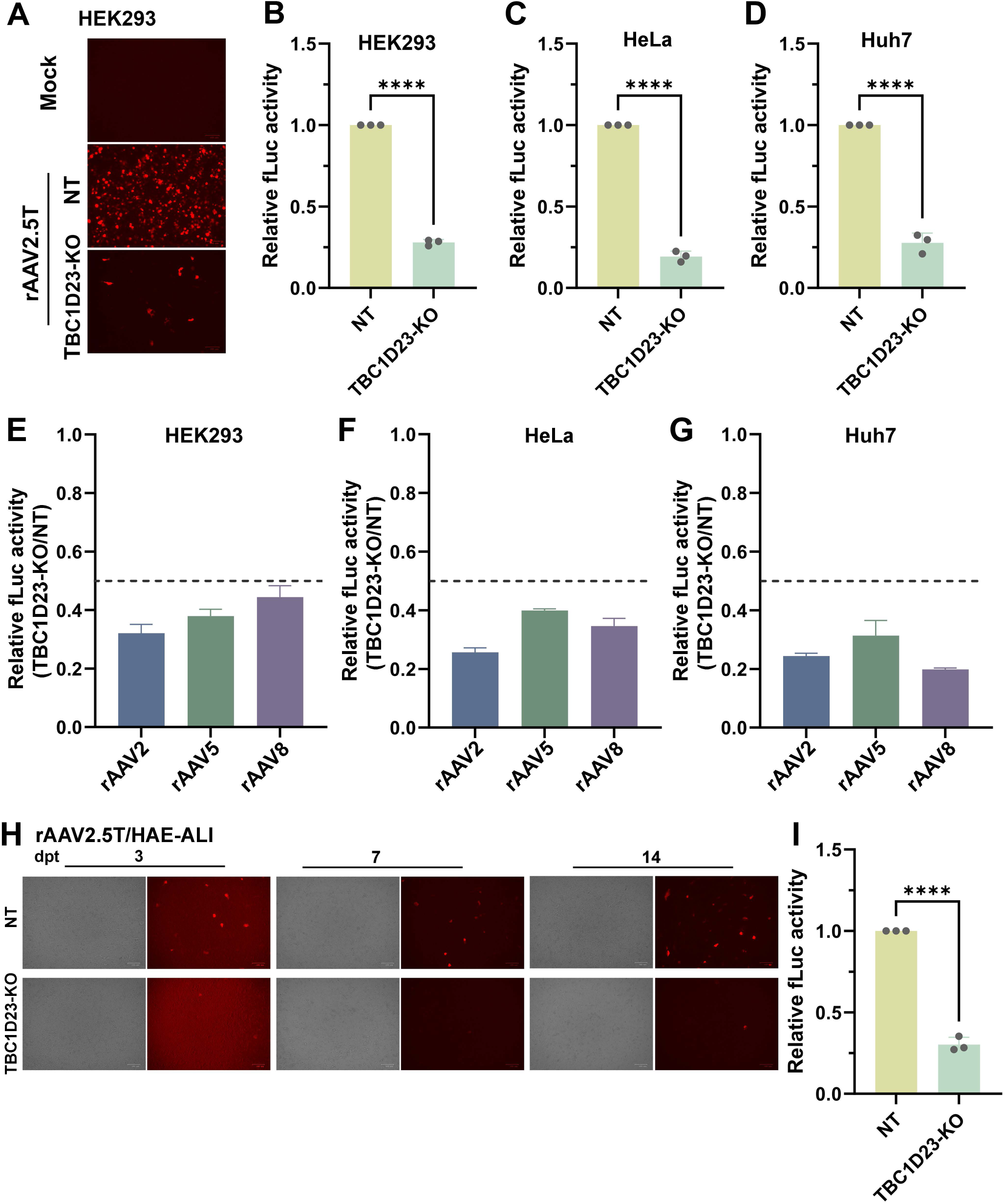
TBC1D23 is required for efficient rAAV transduction in cell lines and polarized airway epithelium. (A&B) rAAV2.5T transduction of TBC1D23-KO HEK293 cells. (A) mCherry expression. Representative fluorescence images of HEK293 cells transduced with rAAV2.5T in non-targeting control (NT) or TBC1D23-KO cells at an MOI of 20K. Cells were imaged at 2 days post-transduction (dpt). Mock-infected cells are shown as negative control. (B) Quantification of firefly luciferase (fLuc) activity. NT and TBC1D23-KO HEK293 were transduced with rAAV2.5T at an MOI of 20K. At 2 dpt, cells were lysed for fLuc activity. Data are normalized to NT control cells. **(C&D) rAAV2.5T transduction of TBC1D23-KO HeLa and Huh7 cells.** NT and TBC1D23-KO HeLa and Huh7 cells were transduced with rAAV2.5T at an MOI of 20K. At 2 dpt, cells were lysed for fLuc activity. Data are normalized to NT control cells **(E-G) Transduction of AAVR-dependent AAVs in TBC1D23-KO cells.** Transduction of rAAV serotypes (rAAV2, 5, and 8) in TBC1D23-KO HEK293, HeLa, and Huh7 cells at an MOI of 20K was compared with NT controls at 2 dpt. Data are normalized to NT cells for each serotype. The dashed line indicates 50% transduction efficiency relative to the control. Data are normalized to NT cultures. **(H&I) rAAV2.5T transduction of HAE-ALI cultures.** (H) Representative fluorescence and brightfield images of polarized HAE-ALI cultures transduced with rAAV2.5T at 3, 7, and 14 dpt, respectively. Reduced reporter expression is observed in TBC1D23-KO ALI cultures compared with NT controls over time. (I) Quantification of fLuc activity in NT and TBC1D23-KO HAE-ALI cultures following rAAV2.5T transduction. Mean ± SD is shown (n = 3 independent ALI cultures). ****P < 0.0001.

TBC1D23 has an established role in retrograde transport from endosomes to the TGN (18), which is aligned with a proposed retrograde transport pathway for AAV trafficking (7). Together with our findings, these observations led us to hypothesize that TBC1D23 interacts with AAVR and functions as a trafficking adaptor that links AAVR to retrograde transport machinery.

### TBC1D23 is essential for rAAV transduction in multiple cell lines and in polarized human airway epithelium

To determine whether TBC1D23 is essential for rAAV transduction in other cells/tissues, we used CRISPR/Cas9 to generate TBC1D23-KO HeLa and Huh7 cell lines. Loss of TBC1D23 protein expression in these cell lines was confirmed by immunoblotting (**Fig. S4**). Cells were then transduced with rAAV2.5T encoding dual mCherry and fLuc reporters. TBC1D23-KO resulted in a profound reduction in transduction efficiency across all three lines, compared with control cell lines treated with non-target sgRNA (NT). Specifically, the luciferase activity was reduced by 72.4%-80.7% (**Fig. 3C&D**). Consistent with these findings, TBC1D23-KO also reduced the transduction of rAAV2, rAAV5, and rAAV8 by 55.6-80.2% in HEK293, HeLa, and Huh7 cells (**Fig. 3E-G**). Notably, all the tested AAV serotypes are known to be AAVR-dependent (9).

To assess the relevance of TBC1D23 in a physiologically relevant model, we next examined the impact of its loss on rAAV2.5T transduction in human airway epithelium polarized at an air–liquid interface (HAE-ALI). These cultures recapitulate the pseudostratified architecture of the human proximal airways (19,20) and are commonly used to test gene delivery by rAAV vectors (21–24). Using lentiviral delivery of the CRISPR/Cas9 components, we generated TBC1D23-deficient ALI cultures with the loss of TBC1D23 confirmed at the protein level (**Fig. S4**). Apical exposure of these cultures to rAAV2.5T resulted in a drastic reduction in transduction in TBC1D23-deficient ALI cultures, compared with NT control cultures, as assessed by both mCherry expression and fLuc activity (**Fig. 3H&I**). These results demonstrate that TBC1D23 is required for rAAV transduction in differentiated HAE, highlighting its relevance for in vivo gene therapy of lung diseases, such as cystic fibrosis (25).

Collectively, these results established TBC1D23 as an essential host factor for efficient rAAV transduction across multiple AAV serotypes and cell types, including polarized HAE. Importantly, the drastic reduction in transduction observed upon TBC1D23 depletion support a critical role for TBC1D23 in AAV post-entry intracellular trafficking.

### AAVR and TBC1D23 interact directly via their C-terminal domains with nanomolar affinity

TBC1D23 is a Golgi-localized protein that functions as a vesicle-tethering adaptor mediating endosome-to–TGN trafficking through its C-terminal pleckstrin homology (PH)-like domain (TBC1D23-C), which lacks canonical phosphoinositide-binding motifs (12–14). Based on this functional distinction, we examined a direct interaction of TBC1D23-C with AAVR-C in vitro. GST-tagged AAVR-C (GST-AAVR-C) and His-tagged C-terminal domain of TBC1D23 (TBC1D23-C^His^) were expressed and purified from *E. Coli* (**Fig. 4A**). GST-AAVR-C efficiently pulled down TBC1D23-C^His^, as shown by Coomassie blue staining and Western blotting visualized using specific antibodies (**Fig. 4A-C**), indicating a direct interaction between TBC1D23-C and AAVR-C. Next, we quantified this interaction using biolayer interferometry (BLI). TBC1D23-C^Flag^ was immobilized on Ni-NTA biosensors, and GST-AAVR-C was used as the analyte. Binding curves yielded an equilibrium dissociation constant (K_D_) of 44.2 nM (**Fig. 4D**). Association and dissociation kinetics were consistent with a stable, high-affinity scaffold– receptor interaction. No measurable binding was observed when GST alone was introduced at similar concentrations (**Fig. 4E**).

**Fig. 4.**
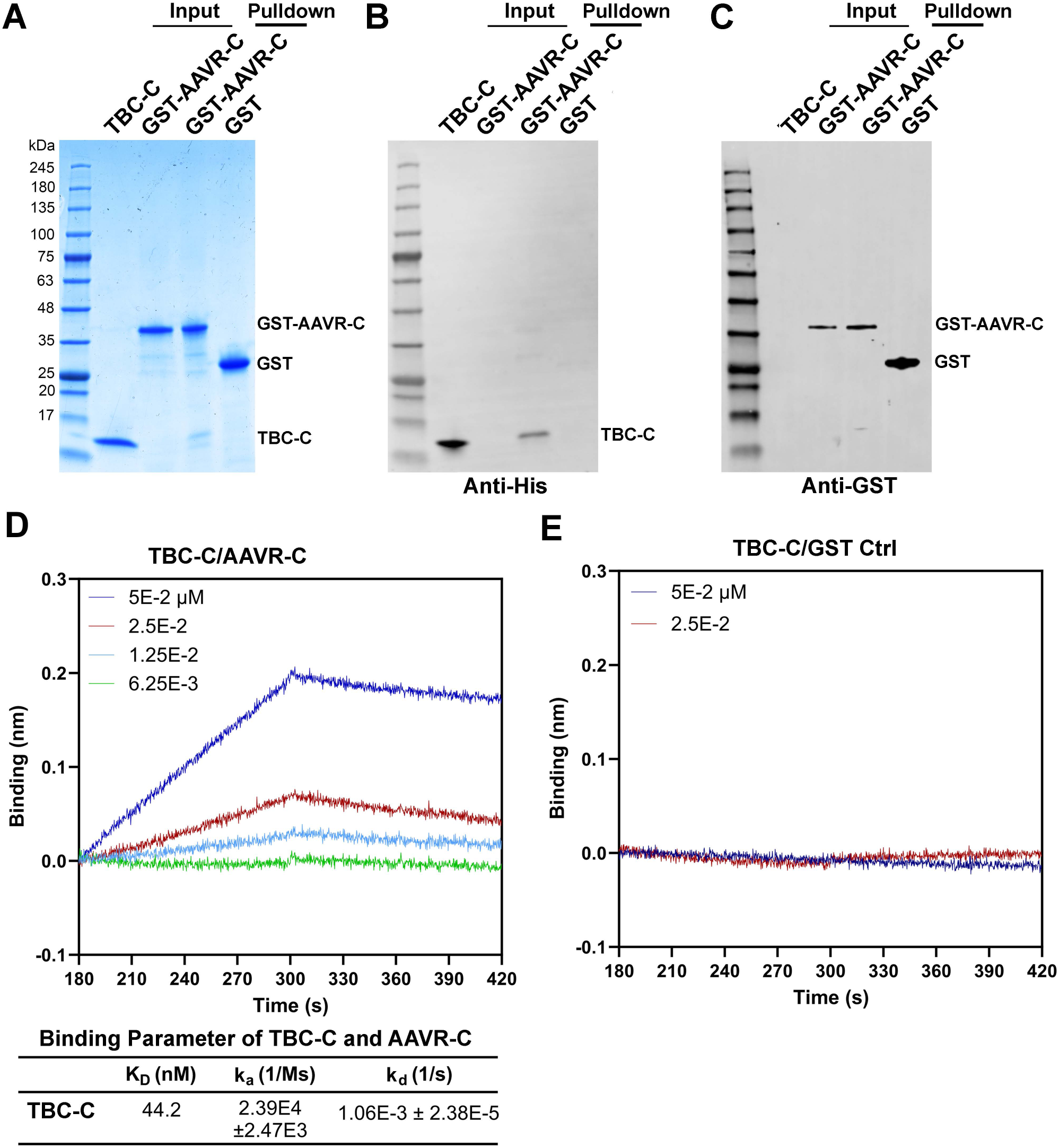
Direct interaction between the AAVR C-terminal cytosolic tail and TBC1D23 C terminus. (A-C) In vitro GST pulldown assay. TBC1D23-C^His^ (TBC-C) and GST-AAVR-C (AAVR-C) were purified for the pulldown assays. “Input” lanes: TBC-C and GST-AAVR-C were loaded alone; “Pulldown” lanes: GST-AAVR-C and GST (control) were used to pull down TBC-C, respectively. Proteins were analyzed on SDS–PAGE by Coomassie Blue staining (A) and immunoblotting with anti-His (B) and anti-GST (C) antibodies, respectively. The identities of the detected bands are indicated. **(D&E) Biolayer interferometry (BLI) analysis.** The TBC1D23-C^His^ (TBC-C) was immobilized to the His biosensors, and GST–AAVR-C (AAVR-C) (D) or GST as a control (E) was added at various concentrations, as shown in the BLI assays. Representative sensorgrams show concentration-dependent binding. Global fitting of the data yields kinetic parameters (k_a_ and k_d_) and an equilibrium dissociation constant (K_D_) of 44.2 nM (D).

Collectively, these data demonstrate a direct and selective interaction between the cytosolic tail of AAVR and the C-terminal scaffold domain of TBC1D23 at a nanomolar binding affinity, supporting a direct receptor–adaptor relationship rather than an indirect association mediated by larger Golgi complexes.

### The acidic residue cluster (ARC) of the AAVR cytosolic tail is the principal determinant for recruitment of TBC1D23

We next sought to define the functional determinants in AAVR-C required to mediate its interaction with the C-terminal domain of TBC1D23. First, we analyzed the sorting signals within AAVR-C (26,27) (**Fig. 5A**) and predicted the structure of AAVR-C with the resolved TBC1D23-C structure (PDB #6JM5) using AlphaFold 3 (15,28) (**Fig. S5A&B**). Structural modeling predicted that the acidic residue cluster (ARC) within AAVR-C closely interacts with TBC1D23-C (**Fig. 5B-D**), but the YXXΦ-like motif did not contribute to their binding (**Fig. S5C&D**). To experimentally prove the function of the ARC motif in the interaction of AAVR-C with TBC1D23-C, we generated an ARC-mutant AAVR-C^mARC^ (M1) and, as a control, a dileucine-mutant AAVR-C^mLL^ (M2) (**Fig. 5A**). GST-tagged mutant AAVR-C proteins were expressed in *E coli* and purified to a comparable purity as GST-AAVR-C (**Fig. 5E**). Using BLI, we found that disruption of the ARC motif (AAVR-C^mARC^) completely abolished the binding between AAVR-C and TBC1D23-C, with no detectable interaction observed at concentrations that readily supported binding of the WT protein (GST-AAVR-C) (**Fig. 5F**). As a control, disruption of C-terminal dileucine motif (AAVR-C^mLL^) retained robust binding with TBC1D23-C (K_D_ = 59.7 nM) (**Fig. 5G**).

**Fig. 5.**
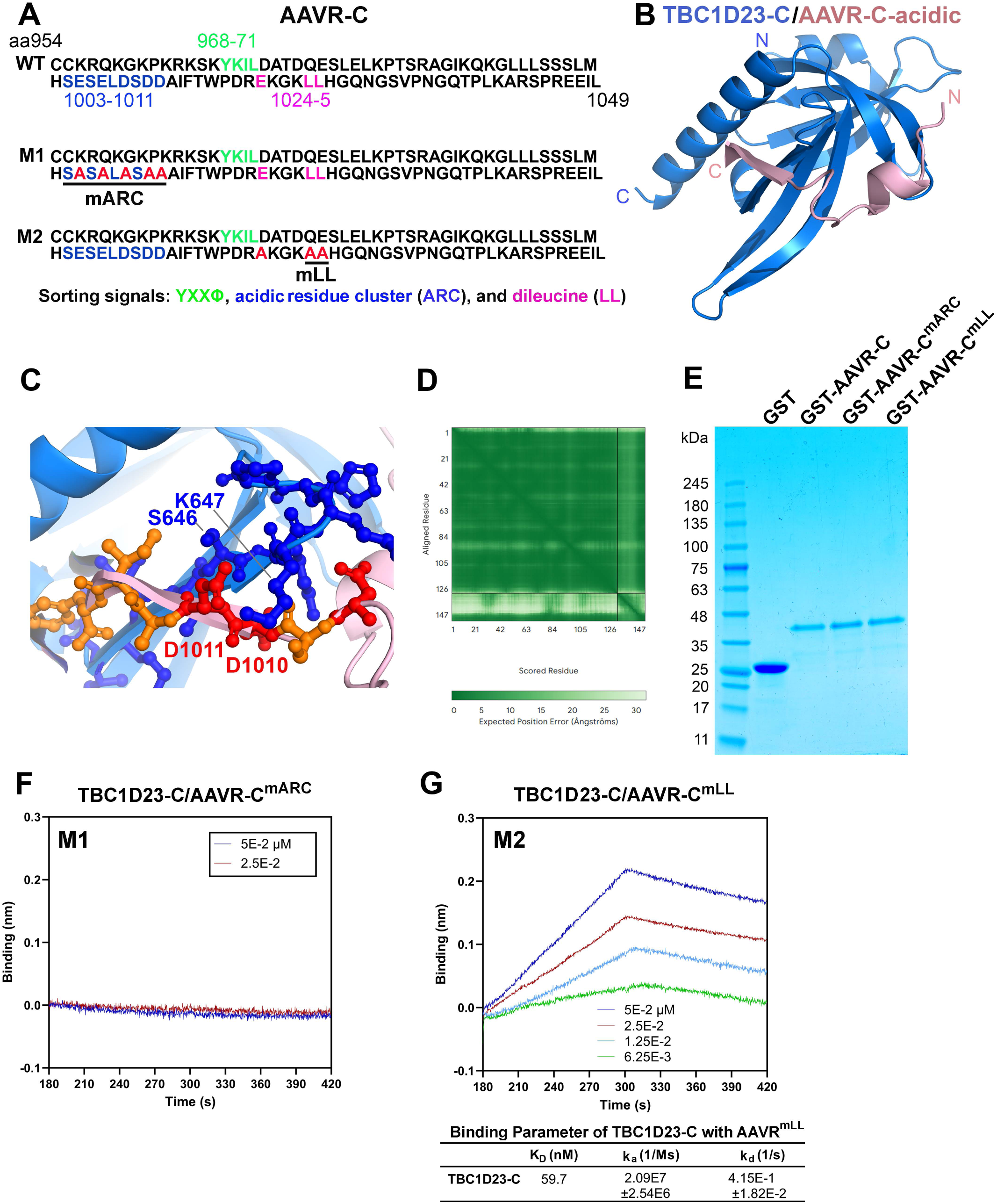
Binding affinity interaction between TBC1D23-C and AAVR-C proteins. (A) Sequences of AAVR-C (aa954-1049) and its mutants. The cytosolic tail of AAVR (AAVR-C) highlights canonical trafficking motifs, including a membrane-proximal YxxΦ motif (green), a central acidic residue cluster (ARC; blue), and a distal dileucine motif (LL; pink). Mutant constructs were generated in which each motif was selectively disrupted (red substitutions) individually. **(B-D) AlphaFold prediction.** (B) AlphaFold-based structural model of TBC1D23-C (aa574-699) (sky blue) in complex with the acidic region of AAVR-C (acidic, aa994-1013) (light purple). The predicted confidence scores for ipTM and pTM are 0.79 and 0.87, respectively. (C) Close-up view of the interface between TBC1D23-C and AAVR-C-acidic, highlighting charged residues within the acidic cluster (red) and basic residues (orange) involved in the interaction. Key acidic residues (D1010 and D1011) are indicated. (D) Predicted aligned error (PAE) plot. The heat map represents the expected positional error (Å) between residue pairs, with lower values indicating higher confidence in the relative orientation of residues. **(E) Protein purification.** Purified recombinant GST, GST-AAVR-C WT, and the mARC and mLL mutants used for in vitro binding assays were analyzed on SDS-PAGE and Coomassie Brilliant Blue–stained. **(F&G) Biolayer interferometry (BLI) analysis.** The interaction between immobilized TBC1D23-C^His^ and GST–AAVR-C bearing mutations was analyzed. (F) BLI sensorgrams showing binding responses of immobilized TBC1D23-C^His^ to GST-AAVR-C^mARC^ at the indicated concentrations. (G) BLI sensorgrams showing binding responses of immobilized TBC1D23-C^His^ to GST-AAVR-C^mLL^ at the indicated concentrations. Global fitting of the data yielded kinetic parameters (k_a_ and k_d_) and an equilibrium dissociation constant (K_D_) of ∼60 nM, indicating that the mLL mutation preserves TBC1D23-C binding to AAVR-C.

Taken together, the high-affinity binding of AAVR-C to TBC1D23-C via its ARC motif suggests that the ARC is the trafficking signal of AAVR and the essential determinant required for the recognition of AAVR by TBC1D23.

### The acidic residue cluster is essential for functional endosome-to-TGN trafficking of the AAV–AAVR complex

To evaluate the functional contribution of the ARC motif to AAV trafficking, AAVR-KO cells were stably transduced with lentiviral vectors expressing codon-optimized AAVR^WT^ and the corresponding ARC or dileucine mutants. Immunoblot analyses confirmed robust overexpression of AAVR^WT^ and mutants (AAVR^mARC^ and AAVR^mLL^) relative to endogenous AAVR expression in WT HEK293 cells (**Fig. 6A**). Complementation with AAVR^WT^ in AAVR-KO HEK293 cells enhanced rAAV transduction by 8-fold, compared with the level in WT HEK293 cells (**Fig. 6B&C**). In contrast, expression of AAVR^mARC^ failed to rescue AAV transduction, phenocopying AAVR-KO cells, whereas the dileucine motif mutant (AAVR^mLL^) rescued rAAV2 transduction as efficiently as AAVR^WT^ (**Fig. 6B&C**).

**Fig. 6.**
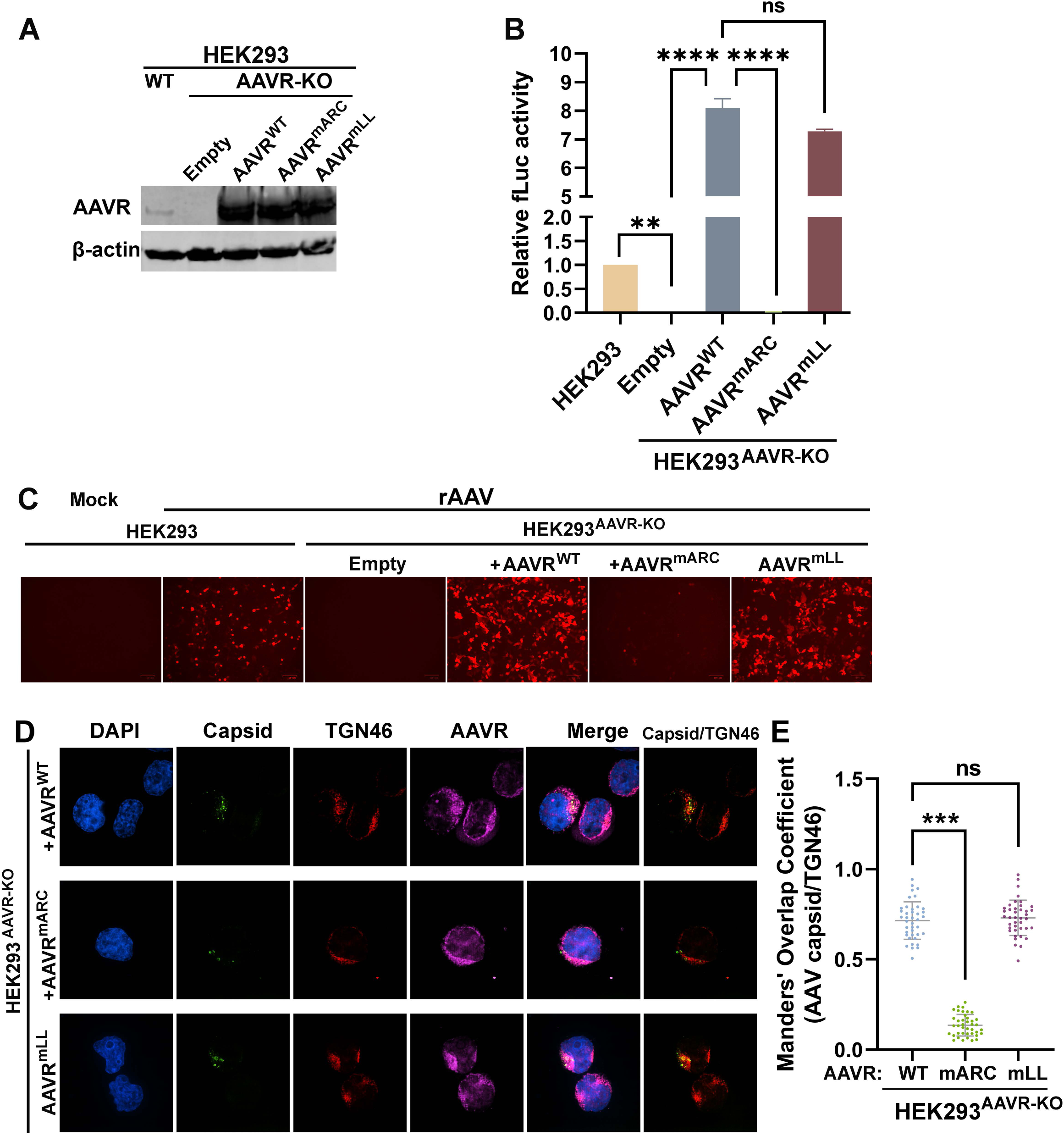
The AAVR cytosolic acidic cluster (ARC) is required for productive rAAV transduction and rAAV vector localization at TGN. (A) Western blotting. AAVR expressions in WT, AAVR-KO (Empty), and AAVR-complemented HEK293^AAVR-KO^ cells were analyzed by Western blotting using anti-AAVR. β-actin was probed as a loading control. **(B&C) rAAV2.5T transduction.** WT, AAVR-KO (Empty), AAVR, AAVR^mARC^, AAVR^mLL^-complemented HEK293^AAVR-KO^ cells were transduced with rAAV2.5T at an MOI of 10K. (B) fLuc assay. At 2 dpt, transduction was measured by luciferase activity. Data are normalized to WT control cells. (C) mCherry expression. At 2 dpt, cells were imaged for mCherry fluorescence expression. Representative images are shown. **(D&E) AAV TGN localization.** Confocal microscopy shows colocalization of AAV2.5T capsid with TGN46, and AAVR (stained with anti-AAVR) in the cytoplasm of HEK293^AAVR-KO^ cells complemented with AAVR and its mutants (D). Images were taken under a confocal microscope (CSU-W1 SoRa; Nikon) at x60 magnification. Colocalization of AAV capsid with TGN46 was quantified (E). Manders’ Overlap Coefficient was calculated for AAV capsid colocalization with the TGN46 in single cells (n = 40 per group) using ImageJ (Fiji). Each dot represents one cell. Bars show mean ± SD. **P < 0.01; ***P < 0.001; ****P < 0.0001; and ns, P > 0.05.

Confocal microscopy revealed that AAVR^WT^ supported efficient delivery of AAV capsids to the TGN, as shown by robust colocalization with the TGN46 marker (**Fig. 6D&E**). Whereas AAVR^mARC^ prevented the accumulation of AAV capsids in the TGN compartments. Cells expressing AAVR^mLL^ exhibited trafficking patterns indistinguishable from those expressing AAVR^WT^ (**Fig. 6D&E**).

Together, these data demonstrate that the AAVR ARC motif is indispensable for endosome-to-TGN trafficking of the AAV-AAVR complex and for productive rAAV transduction.

### The acidic residue cluster motif plays an important role in rAAV nuclear import but not vector entry

We next dissected the contribution of the AAVR-C motifs to vector internalization and nuclear import of various rAAV vectors through evaluation of the functions of the mutants relative to WT (**Fig. 7A**). HEK293^AAVR-KO^ cells complemented with AAVR WT, mARC-, or mLL-mutant were transduced with rAAV2.5T. At 2 hpt, vector internalization assay revealed no significant differences in internalized vector copies among WT-and mutant AAVR-expressing cells, indicating that neither mutation compromised vector entry (**Fig. 7B**). We further assessed nuclear import using subcellular fractionation to quantify vector distribution between cytoplasmic and nuclear compartments. Disruption of the ARC motif significantly reduced nuclear accumulation of vector genomes, whereas the dileucine mutation behaved similarly to AAVR^WT^ (**Fig. 7C**, AAVR^mARC^ vs. AAVR^mLL^), suggesting that the ARC motif primarily controls a post-internalization step essential for vector nuclear import. In addition, we evaluated rAAV5 and rAAV8 transduction in HEK293^AAVR-KO^ cells complemented with AAVR^WT^, mARC-or mLL-mutant. Consistent with the AAV2.5T results, both rAAV5 and rAAV8 transduction were reduced by 98-99% in HEK293^AAVR-KO+mARC^ cells compared with HEK293^AAVR-KO+WT^ cells (**Fig. 7D&G**). Nuclear import of both vectors was significantly decreased in HEK293^AAVR-KO+mARC^ cells, but not HEK293^AAVR-KO+mLL^ and HEK293^AAVR-KO+WT^ cells (**Fig. 7F&I**), while vector internalization remained unaffected in all cases (**Fig. 7E&H**).

**Fig. 7.**
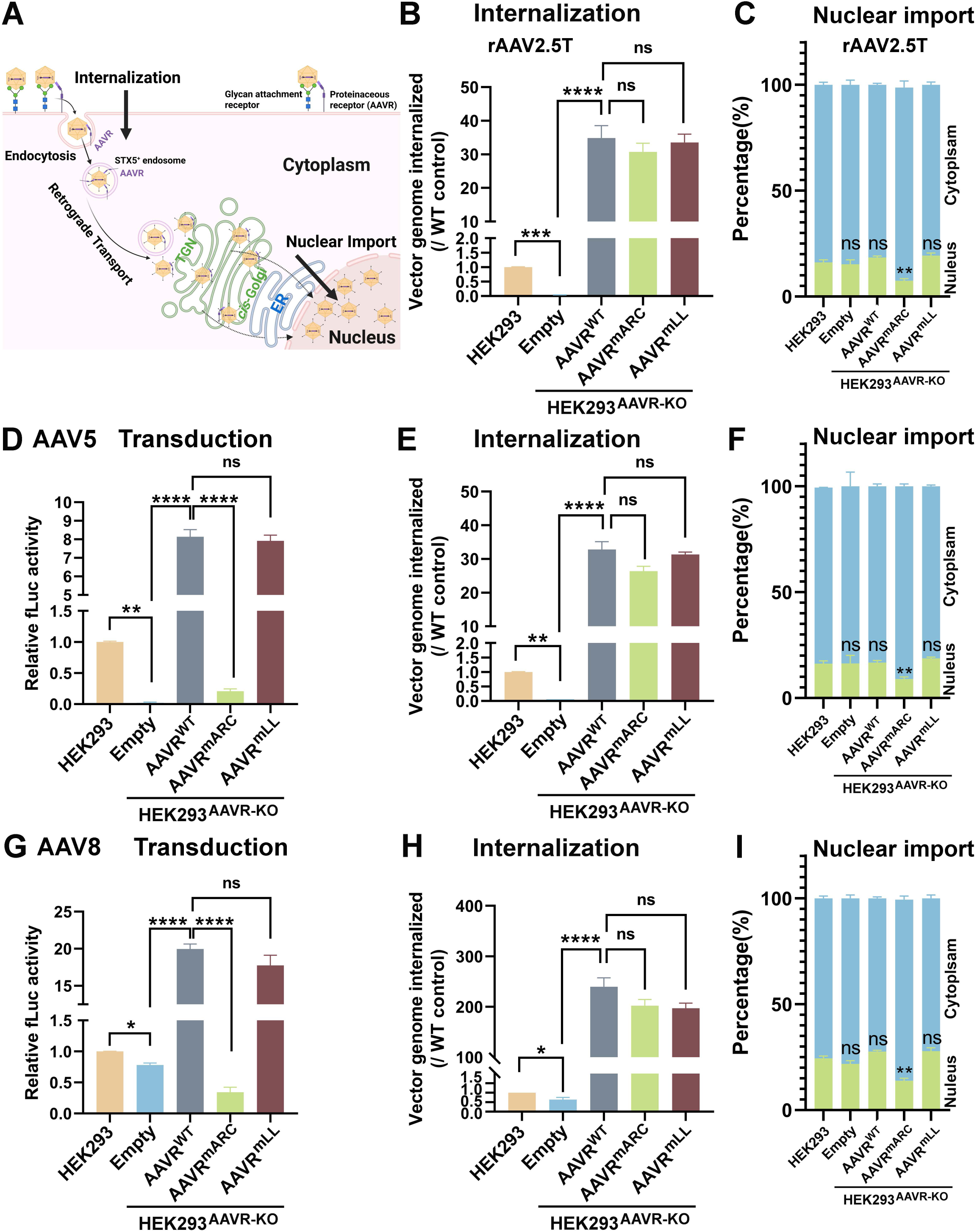
The AAVR cytosolic acidic residue cluster is required for AAV vector nuclear import but not internalization. (A) A diagram of the AAV capsid intracellular trafficking and the analyses. The sequential steps of AAV internalization, intracellular trafficking, and nuclear import are illustrated (7). At the cell surface, AAV particles first engage the primary glycan receptor and AAVR, followed by internalization, which is experimentally assessed using trypsinization to remove non-internalized virions. After endocytosis, the AAV-AAVR complex traffics through STX5⁺ early endosomes and then moves along the retrograde transport pathway toward the TGN and associated Golgi compartments. From the TGN, capsids are either further processed for retrograde transport or released into the cytoplasm and subsequently delivered to the nuclear membrane, where nuclear import occurs. Fractionation of cytoplasmic and nuclear compartments allows quantification of vectors that successfully reach the nucleus. **(B&C) rAAV2.5T vector internalization and nuclear import.** WT, AAVR-KO, and AAVR-complemented HEK293^AAVR-KO^ cells, as indicated, were transduced with rAAV2.5T at an MOI of 5K. At 2 hpt, vector internalization (B) and nuclear import assay (C) were carried out. Vector genomes internalized relative to WT HEK293 cells were calculated (B). % of vectors in the cytoplasm and nucleus are expressed (C). **(D-I) AAV transduction, internalization, and nuclear import measurement.** WT, AAVR-KO, AAVR-complemented HEK293^AAVR-KO^, as indicated, were transduced with rAAV5 at an MOI of 5K (D-E) or AAV8 (G-I) at an MOI of 5K. At 2 dpt, transduction efficiency was assessed by fLuc activity assay (D&G). At 2 hpt, vector internalization (E&H) and nuclear import assays (F&I) were carried out. For vector transduction and internalization assays, data are normalized to WT HEK293 cell controls. For the nuclear import assay, % of vectors in the cytoplasm and nucleus are expressed. All data are shown as mean ± SD (n = 3). *P < 0.05; **P < 0.01; ***P < 0.001; ****P < 0.0001; and ns, P > 0.05.

Notably, AAV8 transduction of HEK293^AAVR-KO^ cells was decreased only by 15% (**Fig. 7G**), in contrast to the >95% reduction observed in HeLa^AAVR-KO^ cells (9). However, upon AAVR overexpression, AAV8 transduction became strongly AAVR-dependent, as expression of the ARC mutant (AAVR^mARC^) reduced transduction by >90%, compared to the expression of AAVR^WT^. Again, this reduction correlated with the impaired vector nuclear import but was not due to altered vector internalization (**Fig. 7H&I**).

Taken together, these results indicate that the ARC motif of AAVR functions in AAV intracellular trafficking downstream of vector internalization and upstream of nuclear import.

### TBC1D23 is required for endosome-to-Golgi trafficking of the AAV capsid-AAVR complex

We next examined how TBC1D23 affects AAV intracellular trafficking. To this end, NT control and TBC1D23-KO HEK293 cells were transduced with AAV5 and analyzed by confocal microscopy after fixation at 6 hpt. In NT control cells, incoming AAV capsids accumulated in perinuclear regions that co-stained with the TGN marker TGN46 and with AAVR (**Fig. 8A**, NT). However, this trafficking pattern was disrupted in TBC1D23-KO cells. Capsids remained in peripheral puncta but showed little overlap with TGN46 (**Fig. 8A**, TBC1D23-KO). Quantitative colocalization analysis confirmed a significant decrease in AAV capsid-TGN overlap in the absence of TBC1D23 (**Fig. 8B**). Because loss of TBC1D23 has been reported to affect TGN46 localization (12), we instead used GM130, a *cis*-Golgi marker that is independent of TBC1D23-mediated retrograde trafficking, to assess Golgi-associated localization of the AAV-AAVR complex. Consistent with the above findings, AAV5 capsid–AAVR complexes remained in peripheral puncta and exhibited little colocalization with GM130 in TBC1D23-KO cells (**Fig. S6A&B**, TBC1D23-KO).

**Fig. 8.**
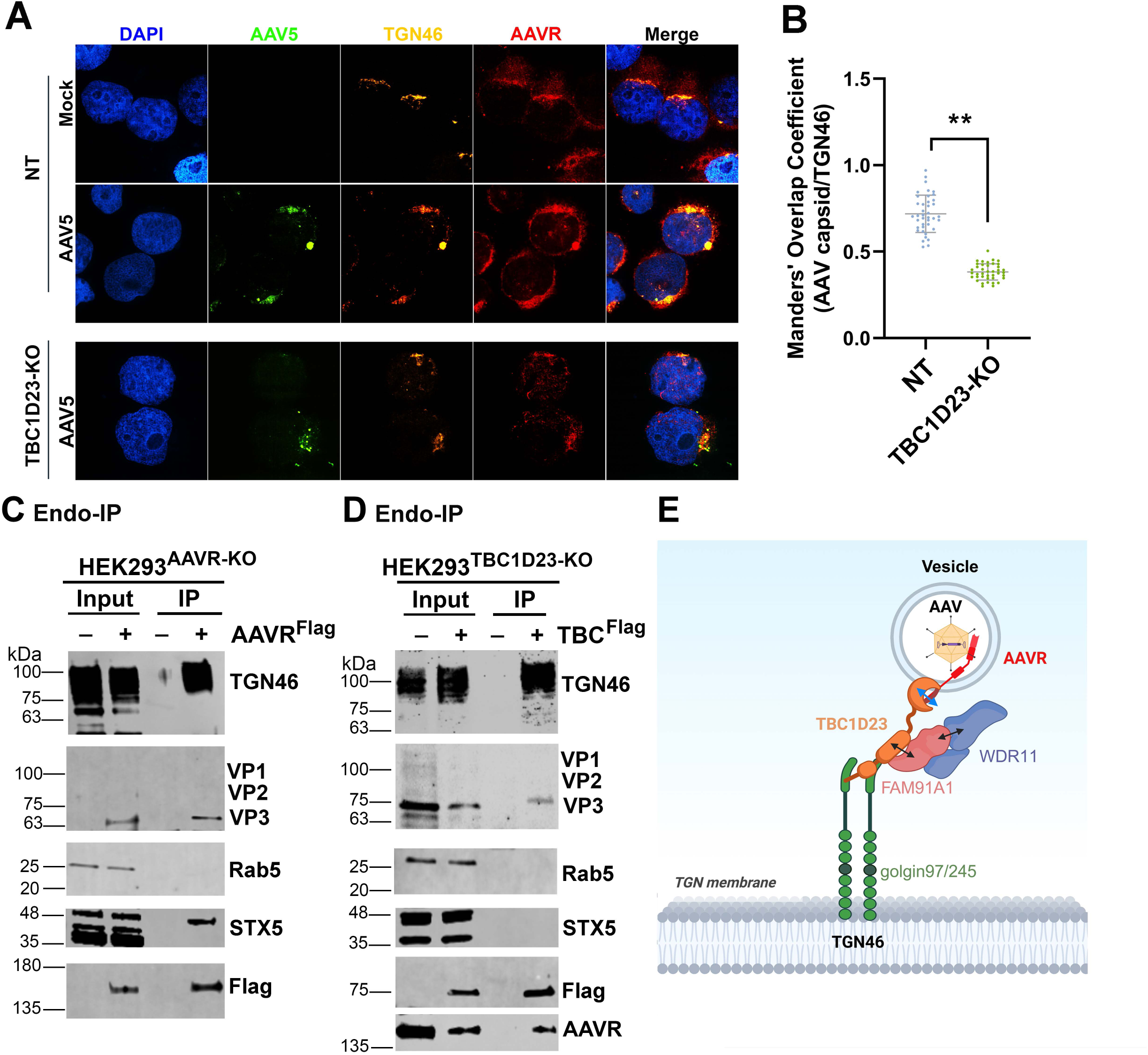
TBC1D23 promotes trafficking of the AAVR–AAV complex from the endosome to the TGN. (A&B) TBC1D23-KO disrupts the localization of AAV capsids to the TGN. NT and TBC1D23-KO HEK293 cells were transduced with rAAV5 at an MOI of 50K. At 6 hpt, cells were stained for AAV capsid, the TGN marker TGN46, and AAVR, and imaged under a confocal microscope (CSU-W1 SoRa; Nikon) at x 60 magnitude (A). Manders’ Overlap Coefficient was calculated for AAV colocalization with the indicated markers in single cells (n = 40 per group) using ImageJ (Fiji). Each dot represents one cell; bars show mean ± SD (B). **P < 0.01. **(C&D) Endosomal immunoprecipitation (Endo-IP) assays**. (C) HEK293^AAVR-KO^ (-) and HEK293^AAVR-KO/WT^ (+) cells that are reconstituted with WT AAVR^FLAG^, or (D) HEK293^TBC1D23-KO^ (-) and HEK293^TBC1D23-KO/WT^ (+) cells reconstituted with WT TBC1D23^FLAG^ were transduced with rAAV5 at an MOI of 50K. At 2 hpt, cells were washed, lysed under detergent-free conditions by Dounce homogenization, clarified, and Flag-tagged vesicles were immunoprecipitated using anti-Flag M2 magnetic beads. Inputs (10%) and IP eluates were analyzed by immunoblotting for the indicated markers (TGN46, STX5, Rab5, and AAVR) and AAV capsid proteins (VP1/VP2/VP3). VP3 was readily detected, whereas VP1/VP2 were at or below the detection limit, consistent with the lower stoichiometry of VP1/VP2 relative to VP3 and limited capsid abundance in Endo-IP samples. Input: 10% of the cell lysates used for IP. Protein ladders are indicated to the left. **(E) A schematic model illustrating the role of TBC1D23 in AAVR-mediated endosome-to-TGN trafficking.** Schematic illustration depicts how internalized AAV particles, bound to the proteinaceous receptor AAVR, are delivered to the TGN. Following endocytosis, AAV-containing endosomal vesicles engage TBC1D23, which acts as a vesicle-tethering adaptor by linking the cytosolic tail of AAVR to the Golgi-anchored WDR11–FAM91A1 complex. This tethering complex is stabilized at the TGN membrane through interactions with golgin-97/245, thereby promoting vesicle capture, docking, and convergence of AAV-containing carriers at the TGN prior to downstream trafficking steps. The model incorporates established interactions reported previously for TBC1D23, WDR11, FAM91A1, and golgins (12,13,31).

To determine if this defect was driven by altered AAVR localization or abundance, we examined endogenous AAVR distribution in NT and TBC1D23-KO HEK293 cells in the absence of AAV transduction. Confocal microscopy showed that endogenous AAVR remained detectable in TBC1D23-KO cells but exhibited reduced colocalization with GM130 compared with NT cells (**Fig. S6C**), indicating impaired Golgi-associated localization. In contrast, immunoblot analysis revealed comparable AAVR protein levels between NT and TBC1D23-KO cells (**Fig. S6D**). Together, these findings demonstrated that TBC1D23 regulates the subcellular localization of AAVR rather than its stability or abundance, supporting a role of TBC1D23 in transporting AAV capsid-AAVR complexes to TGN, thereby facilitating efficient vector nuclear import (**Fig. 7**). Moreover, overexpression of TBC1D23 increased AAV capsid nuclear localization (**Fig. S7A&B**) and enhanced transduction (**Fig. S7C**), further supporting the role of TBC1D23 in transporting AAV capsid to TGN, increasing vector nuclear import and transduction. These findings indicate that TBC1D23 is required for retrograde trafficking of AAV capsids from peripheral endosomes to the TGN.

To further define the vesicular compartments involved, we carried out an endosome-immunoprecipitation assay (endo-IP) for the association of AAV capsid-AAVR with TGN and other vesicles. HEK293^AAVR-KO^ cells complemented with Flag-tagged AAVR^WT^ (HEK293^AAVR-KO+WT^) were transduced with AAV5. The cells were harvested at 2 hpt for anti-Flag endo-IP to pull down AAVR-associated membrane vesicles. The results showed pulldown of AAV VP, TGN46, and STX5, but not Rab5 (**Fig. 8C**), suggesting that AAVR is associated with TGN46^+^ TGN and STX5^+^ endosomes, but not Rab5^+^ early endosome. In parallel, another endo-IP, which used anti-Flag to pull down Flag-tagged TBC1D23-associated membrane vesicles, was performed in AAV-transduced HEK293^TBC1D23-KO+WT^ cells, which were HEK293^TBC1D23-KO^ cells overexpressed with WT TBC1D23^Flag^. The results showed that AAV VP, TGN46, and AAVR, but not STX5 and Rab5, were pulled down (**Fig. 8D**). As a negative control, anti-Flag beads did not pull down those proteins from Flag-GFP-expressing HEK293 cells (**Fig. S8**). These results demonstrated a close association of the AAV capsid-AAVR complex with TGN and STX5^+^ endosomes, and the capsid-TBC1D23 association with TGN, but not with STX5^+^ endosomes.

Together, these results confirmed that TBC1D23 regulates the subcellular localization of AAVR and plays an important role in transporting the AAV-AAVR complex from the STX5^+^ endosome to the TGN, a retrograde transport pathway that has been previously suggested as a major intracellular trafficking pathway of AAV vectors (10,29).

## Discussion

This study reveals a previously unrecognized role for the vesicle tethering factor TBC1D23 in mediating retrograde transport of AAV. Through a combination of targeted gene KO and complementation, biochemical interaction assays, endosome immunoprecipitation, and confocal imaging-based colocalization analyses, we defined a mechanistic link between AAVR and TBC1D23 during AAV intracellular trafficking, where TBC1D23 is a key player in the endosome-to-TGN transport of the AAV-AAVR complex. Our data demonstrate that the cytoplasmic C-terminal tail of AAVR directly binds the C-terminal scaffold domain of TBC1D23 with nanomolar affinity, positioning TBC1D23 as a receptor-proximal adaptor that couples AAVR-containing endosomal vesicles to the TGN tethering machinery. Notably, this interaction is dispensable for AAV internalization, suggesting that following internalization, AAVR constitutively associates with the retrograde transport pathway, enabling AAV vectors to exploit a pre-existing receptor trafficking circuit to access the TGN.

TBC1D23 has been reported to directly interact with FAM91A1, which cooperates in regulating endosome-to-TGN trafficking of AAVR (12,13,30); however, the underlying mechanism remains unclear. FAM91A1 itself directly interacts with WDR11 (31), and the WDR11-FAM91A1 complex functions downstream of the clathrin-AP-1 machinery to facilitate cargo transport from endosomes to the TGN (32,33). Consistent with this pathway, we observed a close association of incoming AAV capsids with WDR11 at TGN (**Fig. S9A**), and disruption of either WDR11 or FAM91A1 significantly impaired AAV transduction (**Fig. S9B&C**). In addition, although the WDR11 complex is known to facilitate the transport of acidic residue cluster (ARC)-containing cargoes from endosomes to the TGN (32), most notably through direct interaction with an ARC-containing cargo protein, cation-independent mannose-6-phosphate receptor (Ci-MPR) (31). Ci-MPR is the prototypical receptor that mediates trafficking from the plasma membrane to the TGN (34). However, our data indicate that AAVR’s engagement in this pathway appears mediated by TBC1D23. Specifically, in TBC1D23-KO cells, AAVR-C did not interact with WDR11, as shown by co-IP (**Fig. S9D**). Loss of TBC1D23 or mutation of the ARC motif in AAVR selectively blocked AAV capsid delivery to the TGN, leading to accumulation of capsids in peripheral endosomal compartments. These findings align with previous structural and genetic studies of TBC1D23 in other cargo systems (13), underscoring its conserved role in bridging endosomal vesicles to TGN.

Based on these observations, we proposed a model in which the WDR11–FAM91A1 complex cooperates with TBC1D23 to mediate AAVR-dependent retrograde transport of AAV capsids from endosomal compartments to TGN (**Fig. 8E**). Following capsid binding to AAVR through its PKD domains at the plasma membrane, AAV-AAVR complex is endocytosed through the clathrin-independent carriers/GPI-enriched early endosomal compartments (CLIC/GEEC) pathway to STX5^+^ endosome (35), which transports the complex into the TGN and down to the *cis*-Golgi (**Fig. 7A**). The cytoplasmic tail of AAVR is likely extended into the cytosol from the endosomes to recruit TBC1D23 through its C-terminal acidic residue cluster. TBC1D23 then acts as a molecular bridge, connecting the AAVR–AAV complex on the vesicle to the WDR11–FAM91A1 complex, which stabilizes the tether and links it to TGN-resident golgins, such as golgin-97 and golgin-245 (13). Together, these components form a multi-protein tether that captures AAV-containing vesicles, promotes their retrograde transport, and facilitates docking at the TGN membrane. In this model, TBC1D23 serves as the central adaptor, physically coupling AAVR to TGN golgins, FAM91A1, and WDR11 through its N terminal domain ( aa1-331) (14), middle unstructured domain (aa514-538) (13), and C-terminal PH-like domain (aa559-684), respectively, thereby ensuring efficient delivery of AAV to the perinuclear Golgi compartment—a prerequisite for AAV nuclear import and productive transduction. In the TGN where GPR108 (G Protein-Coupled Receptor 108) localizes, the VP1 unique region (VP1u) extended from the surface of AAV capsids likely interacts with GPR108 and mediates the trafficking of AAV capsids from the TGN toward *cis*-Golgi and into the nucleus. GPR108 has been identified as a conserved essential factor for transduction of rAAV2 but not of rAAV5, which is due to the VP1u difference (36,37).

Interestingly, while TBC1D23-KO reduced transduction of multiple AAVR-dependent AAV serotype vectors by ∼75% (**Fig. 3**), mutation of the AAVR ARC motif that disrupts the interaction of AAVR-C with TBC1D23-C reduced AAV transduction by more than 95% (**Fig. 6&7**). This disparity suggests that the ARC motif of AAVR likely interacts with additional sorting adaptors that contribute to AAV retrograde transport to the TGN, warranting further investigation. In addition, we observed that AAV8 transduction is less dependent on AAVR in HEK293 cells than in HeLa cells (9). It has been reported that AAV8 can use an alternate receptor, CPD, for transduction (38). Both CPD and AAVR are cargo-sorting transmembrane proteins associated with endosome-to-TGN transport pathways. TBC1D23 interacts with CPD via its C-terminal domain, which is required for CPD endosome-to-Golgi trafficking (14). Consistent with this, we observed a dependence of rAAV8 transduction on TBC1D23 in TBC1D23-KO cells. However, upon AAVR overexpression, rAAV8 transduction becomes largely AAVR-dependent. We speculate that the AAVR expression levels and the AAVR-AAV8 capsid affinity together determine the receptor usage and trafficking dependence. Distinct AAV serotypes may explore related endosome-to-Golgi trafficking machinery through different host adaptors or cargo receptors.

Collectively, our results extend the functional map of AAVR beyond capsid binding by identifying a downstream molecular interface that is essential for productive rAAV transduction. This work fills a key knowledge gap in AAV receptor biology by elucidating how receptor-bound capsids traverse the complex endosomal network to reach processing compartments. Our findings support a model in which AAV exploits multiple sequential trafficking factors rather than redundant pathways. The AAVR-TBC1D23 interaction appears to function primarily during endosome-to-TGN transport, whereas GPR108 likely acts downstream at the TGN to facilitate subsequent trafficking events. Likewise, CPD may represent a serotype-specific cargo receptor that utilizes overlapping retrograde trafficking machinery. Thus, these factors likely participate in intersecting but non-identical trafficking networks. The direct interaction of AAVR–TBC1D23 positions TBC1D23 at a mechanistic bottleneck in the AAV retrograde transport from endosomes to TGN, making TBC1D23 a potential target for engineering strategies aimed at improving vector tropism or modulating host-vector interaction to enhance rAAV transduction efficiency. From a translational perspective, the TBC1D23-AAVR binding interface could be targeted pharmaceutically to enhance retrograde trafficking, potentially boosting gene transfer efficiency and reducing vector dose in gene therapy applications. Thus, structural characterization of the AAVR–TBC1D23 complex is essential to guide rational design of small-molecule compounds that enhance AAV retrograde trafficking.

In summary, our findings uncover a critical link between the multi-serotype AAVR and the host retrograde trafficking machinery, and establish AAVR as both a viral entry receptor and a cargo-sorting receptor whose cytoplasmic tail actively engages the retrograde trafficking machinery through TBC1D23. This work not only clarifies a missing step in the AAV entry pathway but also identifies a host–vector interface with significant therapeutic potential.

## Materials and Methods

### Cell lines and polarized airway epithelial cultures

HEK293 (#CRL-1573; ATCC), HEK293FT (#R70007, ThermoFisher), HeLa (#CCL-2; ATCC), Huh-7 cells (#CVCL_0336, Cell Lines Service LLC, Sioux Falls, SD), and HEK293^AAVR-KO^ cells (9) were grown in Dulbecco’s Modified Eagle Medium (DMEM; #SH30022.01, Cytiva, Marlborough, MA) supplemented with 10% fetal bovine serum (FBS; #0926, MilliporeSigma, St Louis, MO) and 100 units/mL penicillin-streptomycin (PS) in a humidified incubator with 5% CO_2_ at 37°C.

### Human airway epithelial cells

CuFi-8 cells are human primary tracheal airway epithelial cells derived from a cystic fibrosis patient and immortalized by the expression of human telomerase reverse transcriptase and human papillomavirus E6/E7 oncogenes (39). CuFi-8 cells were cultured on collagen-coated flasks in PneumaCult-Ex Plus medium (#05040; StemCell Technologies, Vancouver, BC).

### Human airway epithelium (HAE) cultured at an air-liquid interface (ALI)

Proliferating CuFi-8 cells were dissociated from flasks and loaded onto Transwell permeable supports (#3470; Costar, Corning, NY) at a density of 1.5 × 10^5^ cells per insert in PneumaCult-Ex Plus medium (40). At 2 – 3 days after seeding, the media were replaced with PneumaCult-ALI medium (#05001; StemCell). The cells were then polarized in PneumaCult-ALI medium at an ALI for 3 – 4 weeks (40). The maturation of the polarized ALI cultures was determined by measuring transepithelial electrical resistance (TEER) with a Millicell ERS 3.0 Digital Voltohmmeter (MilliporeSigma). HAE-ALI cultures with a TEER value of >1,500 Ω·cm² were used in experiments.

### Plasmid Construction

#### rAAV production plasmids

pAAVRep2-Cap2.5T was constructed with AAV2 VP1u and AAV5 VP2/3 gene (GenBank accession NC _006152), with a single amino acid mutation of A581T (41). pAAVRep2-iCap5 and pAAVRep2-iCap8 were constructed by replacing the PHP.eB VP1 gene with the VP1 gene of AAV5 (42) and AAV8 (43), respectively, in pUCmini-iCAP-PHP.eB (44), a gift from Viviana Gradinaru (Addgene plasmid # 103005). The rAAV transgene plasmid pAVF5tg83fLuc-CMVmCherry(4.6kb) and the adenovirus (Ad) helper plasmid pHelper have been described previously (45).

#### Lentiviral vector

Guide(gRNA)-expressing lentiviral vectors for gene KO were constructed by inserting the targeting sequences of single guide (sg)RNAs into lentiCRISPRv2 puro (#52961). The gRNA sequences are as follows: 5’-TTT CCA CTT AGA TTC AGA CC-3’ and 3’ and 5’-AGA CTC CTA TGC ACT CAA CT-3’ for *TBC1D23*; 5’-CAA CGC CCA CAA CAA GGC GG-3’ for *WDR11*; 5’-GTT GCC GGC CAA CGT GAG AC-3’ for *FAM91A1*. A non-targeting (NT) gRNA-expressing vector has been described previously (46).

#### Mammalian cell expression pLenti-based plasmids

WT AAVR (GenBank: Q8IZA0) and the mutants M1 (AAVR^mARC^) and M2 (AAVR^mLL^), were codon-optimized with a Flag tag at the C-terminus. WT TBC1D23 (GenBank: AAH20955) was codon-optimized with a Flag tag at the C-terminus. They were synthesized at Twist Biosciences (South San Francisco, CA) and cloned into pLentiCMV-Blast-empty (#17486, Addgene) to generate pLenti-AAVR^Flag^, AAVR^mARC^ and AAVR^mLL^ and pLenti-TBC1D23^Flag^. A pLenti-GFP^Flag^ construct was generated and used as a control.

#### Bacterial expression plasmids

The cytoplasmic C-terminal domain of human AAVR (aa953-1049), together with the corresponding mutants (M1 and M2), were codon-optimized and synthesized by Twist Bioscience. They were subcloned into the pGEX-4T-1 vector (Cytivia). For expression of the TBC1D23 C-terminal region, truncated cytoplasmic domains (aa559-684) of human TBC1D23 were cloned into the pET-30a (Novagen), resulting in constructs encoding C-terminal His₆-tagged fusion proteins.

### GST pulldown assay and quantitative mass spectrometry

#### GST pulldown assay

5 μg GST-fused protein was incubated with 2.5 μg His-tagged protein in binding buffer (50 mM Tris-HCl, pH 8.0, 150 mM NaCl, 0.05% NP-40,1mM DTT) for 2 h at 4°C. Beads were washed, and bound proteins were eluted in SDS (Sodium Dodecyl Sulfate) sample buffer and analyzed by Western blotting.

Quantitative mass spectrometry (qMS): qMS was performed at the Taplin Mass Spectrometry Facility, Harvard Medical School. On-bead trypsin digestion was performed, and peptides were analyzed on a Velos Orbitrap Elite ion-trap mass spectrometer (Thermo Fisher) (47). Raw data have been deposited in the MassIVE (Mass Spectrometry Interactive Virtual Environment) repository under dataset identifier MSV000100593. Data were searched against the UniProt human database using MaxQuant. Proteins with ≥ 25 unique peptides, fold change > 2^10^ (GST-AAVR-C vs GST control), and *p* < 0.05 were considered significantly enriched (**Fig. 2A**).

#### Biolayer Interferometry (BLI).h

BLI was performed using Octet RED96e (Sartoris, Bohemia, NY) (48–50). Ni-NTA biosensors (#18-5101, Sartoris) were pre-equilibrated in assay buffer (25 mM Tris-HCl, 150 mM NaCl, pH 7.5, 0.04% Tween 20) for 10 min before immobilizing with His_6_-tagged TBC1D23-C at 60 μg/mL in kinetic buffer for 300 seconds (s) to allow proper binding. A baseline reading was recorded after protein loading in the assay buffer for 60 s to ensure stability. The biosensors were dipped into wells containing different concentrations of AAVR-C (WT or mutants) for 300 s. The binding of AAVR-C to the immobilized TBC1D23-C protein was monitored in real-time. The biosensors were transferred back into the assay buffer, and the dissociation of AAVR-C from TBC1D23-C was recorded for 600 s.

#### Lentivirus production and transduction

Lentiviruses were produced by transfecting HEK293FT cells with sgRNA-expressing lentiCRISPRv2 or pLenti-based plasmids, and two packaging plasmids, psPAX2 and pMD2.G, using PEI MAX, concentrated, and titrated by qPCR Lentivirus Titer Kit (#LV900, Abm, BC, Canada) (46,51). Cells were transduced at a multiplicity of infection (MOI) of 5 transduction units per cell.

#### CRISPR/Cas9 knockout cells

Lentiviral particles were transduced into HEK293, HeLa, Huh-7, or CuFi-8 cells, followed by puromycin selection (2 μg/mL). Gene knockouts were confirmed by Western blotting or Sanger sequencing of the targeting loci.

#### rAAV vector production

rAAV was produced by transfection in Viral Production Cells (VPC) 2.0 (ThermoFisher) with triple plasmids (pAAVRepCap, pAVF5tg83fLuc-CMVmCherry, and Ad pHelper) at equal molar ratios, using AAV-MAX Helper-Free AAV Production System Kit (ThermoFisher). rAAV vectors were purified using two rounds of CsCl gradient ultracentrifugation followed by dialysis against phosphate-buffered saline (PBS, pH 7.4) (23,45). Purified vectors were quantified by quantitative (q)PCR using a transgene-specific probe as DNase-resistant particles (DRP), as previously described (45). rAAV2, rAAV5, and AAV8 vectors that package a CMV-driven fLuc gene were also purchased from AAVnerGene (Rockville, MD).

#### AAV transduction assay

For monolayer cell cultures, the cells were seeded overnight in 48-well plates. rAAV was added to each well at the MOI indicated in the figure legends. For the transduction of ALI cultures (#3470, Costar), 100 µL of rAAV2.5T diluted in Dulbecco’s Phosphate Buffered Saline (D-PBS, pH7.4; #SH30028.03, Cytiva) was added to the apical chamber of the transwell at an MOI of 20K DRP/cell. Subsequently, 0.5 mL of culture media was added to the basolateral chamber. After ∼16 h, the apical inoculum and basolateral media were removed, and the transwell inserts were washed three times with D-PBS. Fresh culture media were then added to the basolateral chamber.

At the indicated days post-transduction (dpt), fLuc activity was quantified using the Luciferase Assay System (#E1483, Promega, Madison, WI) on a Synergy LX Reader (BioTek, Santa Clara, CA). In some cases, the transduced cells were imaged for mCherry expression under ZOE Fluorescent Cell Imager (Bio-Rad).

#### AAV vector internalization and nuclear import assays (46,50)

AAV vector internalization: 2 x 10^6^ cells were incubated with rAAV at an MOI of 5K at 37°C for 2 hours, washed, and treated with trypsin at 0.05% (V/W) and 2 mM EDTA for 10 min to remove cell surface-bound virions. Cells were then detached with trypsin, lysed, and the internalized vector genomes were quantified by qPCR.

AAV vector nuclear import assay: 2 x 10^6^ cells were fractionated using the Subcellular Protein Fractionation Kit (#78840, ThermoFisher) (52), for cytoplasmic and nuclear fractions. Vector genomes in each fraction were quantified by qPCR.

Total DNA was extracted from the cells using the DNeasy Blood & Tissue Kit (Qiagen, Hilden, Germany). Vector genomes were quantified by qPCR using primers specific to the rAAV genome (transgene: *mCherry*) as described previously (45). Standard curves were generated using known amounts of vector DNA to calculate genome copy numbers.

#### Endosomal immunoprecipitation (Endo-IP)

AAVR-KO HEK293 cells were transduced with Lenti-AAVR^Flag^, whereas TBC1D23-KO cells were transduced Lenti-TBC1D23^Flag^ or Lenti-GFP^Flag^ as a control. The cells were selected with Blasticidin at 10 µg/mL. After a few passages, the cells were seeded to ∼80% confluence prior to rAAV transduction. We followed Harper Lab’s Endo-IP method (53,54) to isolate AAVR-associated trafficking complexes during intracellular AAV transport.

Briefly, ∼5 × 10^6^ cells were transduced with rAAV at an MOI of 50K DRP/cell in DMEM for 1 h at 4°C, and 2 h at 37°C to allow vector internalization. Cells were then washed three times with ice-cold D-PBS and placed on ice to halt vesicular trafficking. Cells were lysed on ice in 0.5 ml Endo-IP lysis buffer (25 mM KCl, 100 mM potassium phosphate, pH 7.2, 155 mM NaCl) supplemented with a Protease Inhibitor Cocktail (#S8830, MilliporeSigma). Cell suspensions were disrupted using a 2-mL Dounce homogenizer (B pestle; K885300-002) with 30 strokes on ice to preserve intracellular membrane-associated complexes. Lysates were clarified by centrifugation at 16,000 × g for 10 min at 4°C and pre-cleared with protein G magnetic beads for 30 min at 4°C. Cleared lysates were incubated with 60 µL anti-Flag M2 magnetic beads (#M8823, MilliporeSigma) for 2–4 h at 4°C with gentle rotation. Beads were washed five times with lysis buffer to reduce nonspecific binding. Bound complexes were eluted in 2× Laemmli sample buffer at 95°C for 5 min, resolved by SDS–PAGE (Polyacrylamide Gel Electrophoresis), and analyzed by immunoblotting.

### Co-Immunoprecipitation (Co-IP) assay

HEK293 cells were washed twice with cold PBS and lysed with lysis buffer [50 mM Tris-HCl, pH 8.0, 150 mM NaCl, 1% NP-40, and Protease Inhibitor Cocktail (#S8830, MilliporeSigma)] for co-IP following a previously published method (48).

### Confocal microscopy and colocalization analysis

Cells were fixed with 4% paraformaldehyde, permeabilized with 0.1% Triton X-100, and blocked in 5% bovine serum albumin (BSA) in PBS (55). Primary antibodies (anti-AAV capsid, anti-AAVR, anti-TGN46, and anti-TBC1D23) were applied overnight at 4°C, followed by fluorescently conjugated secondary antibodies. Nuclei were stained with DAPI. Images were captured using a confocal microscope (CSU-W1 SoRa, Nikon, Melville, NY). Pearson’s correlation coefficients were calculated using ImageJ (Fiji) for 40 cells per condition.

### SDS–PAGE and Western Blotting

Protein samples were resolved on 4-20% precast polyacrylamide gels (#4561095, Bio-Rad) and transferred to polyvinylidene difluoride membranes (Millipore). Membranes were blocked in 5% (w/v) non-fat milk dissolved in Tris-Buffered Saline with Tween-20 (TBS-T) and probed with primary antibodies overnight at 4°C, followed by infrared dye-conjugated secondary antibodies. The membrane was imaged on a LI-COR Odyssey F imager (LI-COR Biosciences, Lincoln, NB).

### Recombinant proteins

GST-fused proteins were expressed in *E. coli* BL21(DE3)pLysS and purified with Pierce glutathione agarose (#16100, ThermoFisher Scientific), and His-tagged TBC1D23-C fragments were expressed in *E. coli* and purified using Ni-NTA resin (#30210, Qiagen, Germantown, MD), following previously published methods (56–60).

### siRNA knockdown

Three independent DsiRNAs targeting COPG1, ASAP1, and COPS3 were purchased from Integrated DNA Technologies (IDT). SiRNAs were transfected as pooled mixtures using Lipofectamine RNAiMAX (Thermo Fisher) according to the manufacturer’s instructions. Gene knockdown efficiency was confirmed by immunoblot analysis at 48 h post-transfection.

### Antibodies used in the study

#### Primary antibodies

anti-TBC1D23 (#68409-1-Ig, Proteintech), biotinylated anti-AAVX antibody (#7103522100, ThermoFisher), rabbit anti-TBC1D23 (#17002-1-AP, Proteintech, Rosemont, IL), mouse anti-β-actin (#AC004; ABclonal, Woburn, MA), rabbit monoclonal antibodies against Rab5 (#C8B1, #3547T, Cell Signaling Technology, Danvers, MA), anti-Flag (#F1804, MilliporeSigma), anti-WDR11(A10590, ABclonal), anti-FAM91A1(27738-1-AP, Proteintech), anti-STX5 (#26711-1-AP, Proteintech), mouse anti-AAVR (68717-1-Ig, Proteintech), anti-AAV5 (intact particle) (#610147, clone ADK5, ARP, Waltham, MA), rabbit polyclonal anti-TGN46 (#NBP1-49643, Novus Biologicals, Centennial, CO), sheep anti-TGN46 antibody (GTX74290, GeneTex), rabbit anti-AAV VP (#03-61084, ARP), rabbit anti-AAVR (#21016-1-AP, Proteintech), sheep anti-GM130 (#AF8199, R&D Systems, Minneapolis, MN), rat anti-Flag (#637303, Biolegend, San Diego, CA), goat anti-STX5 (#AF5687, R&D Systems), rabbit anti-ASAP1 (#A302-117A, Bethyl Laboratories), COPS3 (#83822-2-RR, Proteintech), LSG1 (*#*17750-1-AP, Proteintech), PREPL (*#*12478-1-AP, Proteintech), VCP (#10736-1-AP, Proteintech), RPL5 (#29092-1-AP, Proteintech), ASCC3 (#A7960, ABclonal), COPG1(#A10551, ABclonal), and GST(#AE001, ABclonal).

#### Secondary antibodies

Alexa Fluor 488-conjugated Goat anti-Mouse IgG (H+L) cross-absorbed secondary antibody (# A-11001), Alexa Fluor 647-conjugated goat anti-mouse IgG (H+L) cross-absorbed secondary antibody (# A-21235), and Alexa Fluor 594-conjugated goat anti-rabbit IgG (H+L) cross-adsorbed secondary antibody (#A-32754) were purchased from Thermo Fisher Scientific. DyLight 800-conjugated anti-rabbit IgG (#5151S), DyLight 800-conjugated anti-mouse IgG (#5257S), DyLight 680-conjugated anti-rabbit IgG (#5366S), and DyLight 680-conjugated anti-mouse IgG (#5470S) were purchased from Cell Signaling (Danvers, MA). Fluorescein-conjugated streptavidin (SA-5001) was obtained from Vector Laboratories (Burlingame, CA).

## Statistical analysis

All data are presented as mean ± standard deviation (SD) obtained from three independent experiments by using GraphPad Prism 10. Statistical significance (P value) was determined by using an unpaired Student’s t-test for two groups or one-way ANOVA with post-hoc Tukey–Kramer multiple-comparison test among more than two groups. ****P < 0.0001, ***P < 0.001, **P < 0.01, and *P < 0.05 were considered statistically significant, and ns represents not statistically significant.

## Data and materials availability

All data and materials used to evaluate the conclusions in this study are presented in the paper and/or the supplemental material. Raw quantitative mass spectrometry data have been deposited in the MassIVE (Mass Spectrometry Interactive Virtual Environment) repository under dataset identifier MSV000100593 and will be released publicly upon publication.

## Acknowledgments

We thank the members of the Qiu laboratory for their helpful and insightful discussions. We are grateful to Dr. Jan Carette (Stanford University) for providing the AAVR-KO HEK293 cells and HeLa cells.

## Fundings

The study was supported by NIH grants AI166293 (J.Q.), AI180416 (J. Q.), HL174593 (J.Q. and Z.Y.), AI182645 (J.Q. and Z.Y.), and Cystic Fibrosis Foundation YAN23G0 (Z.Y.). The Nikon CSU-W1 SoRa was supported by NIH S10 OD 032207 at the University of Kansas Medical Center. The funder had no role in study design, data collection and interpretation, or the decision to submit the work for publication.

## Notes

### Competing Interest Statement

The authors have declared no competing interest.

