## Supplementary Figures and Legends for "TBC1D23-AAVR Interaction Drives Endosome-to-TGN Trafficking Required for rAAV Transduction"

**This PDF file includes:**

Supplemental Table S1 Legend

Supplemental Fig. S1 to S9 and Legends

**Table S1. List of proteins identified through label-free quantitative mass spectrometry (qMS).**

The table includes protein names (references), gene symbols, annotations, molecular weights (MW), the number of identified peptides (unique reads), and the sum intensities of these peptides. Additionally, it presents the calculated P values and fold-changes between the samples pulled down by GST-AAVR-C and the control GST only.

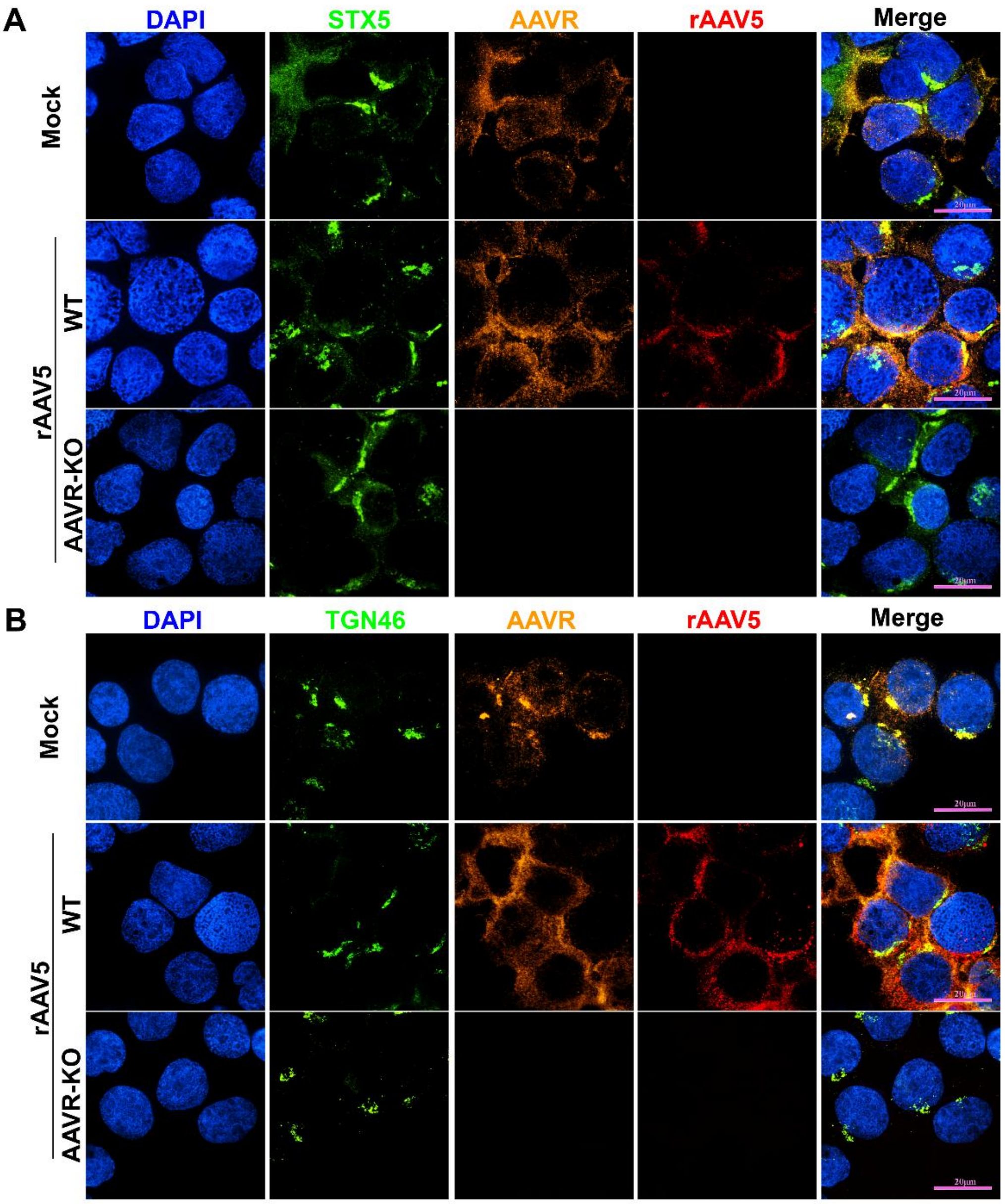


#### Figure S1

**Fig. S1. AAVR colocalizes with Golgi/TGN-associated vesicles during rAAV5 entry.**

Wild-type (WT) or AAVR-knockout (AAVR-KO) HEK293 cells were incubated with rAAV5 at an MOI of 50 K DRP/cell at 4°C for 1 h, followed by incubation at 37°C for various time points post-transduction. **(A) AAV capsid and AAVR colocalize with STX5.** At 4 h post-transduction (hpt), the cells were permeabilized, fixed, and costained with α-STX5 antibody (green), α-AAVR, and α-AAV5 capsid, to label STX5-positive endosomes, AAVR, and AAV5 capsids, respectively**. (B) AAV capsid and AAVR colocalize with TGN46.** At 6 hpt, the cells were permeabilized, fixed, and stained with α-TGN46 antibody, α-AAVR, and α-AAV5 capsid to visualize the trans-Golgi network (TGN), AAVR, and AAV5 capsids, respectively. Nuclei were counterstained with DAPI. Images were acquired using a CSU-W1 SoRa spinning-disk confocal microscope with a 60× objective. Representative images from three independent experiments are shown. Scale bar, 20 μm.

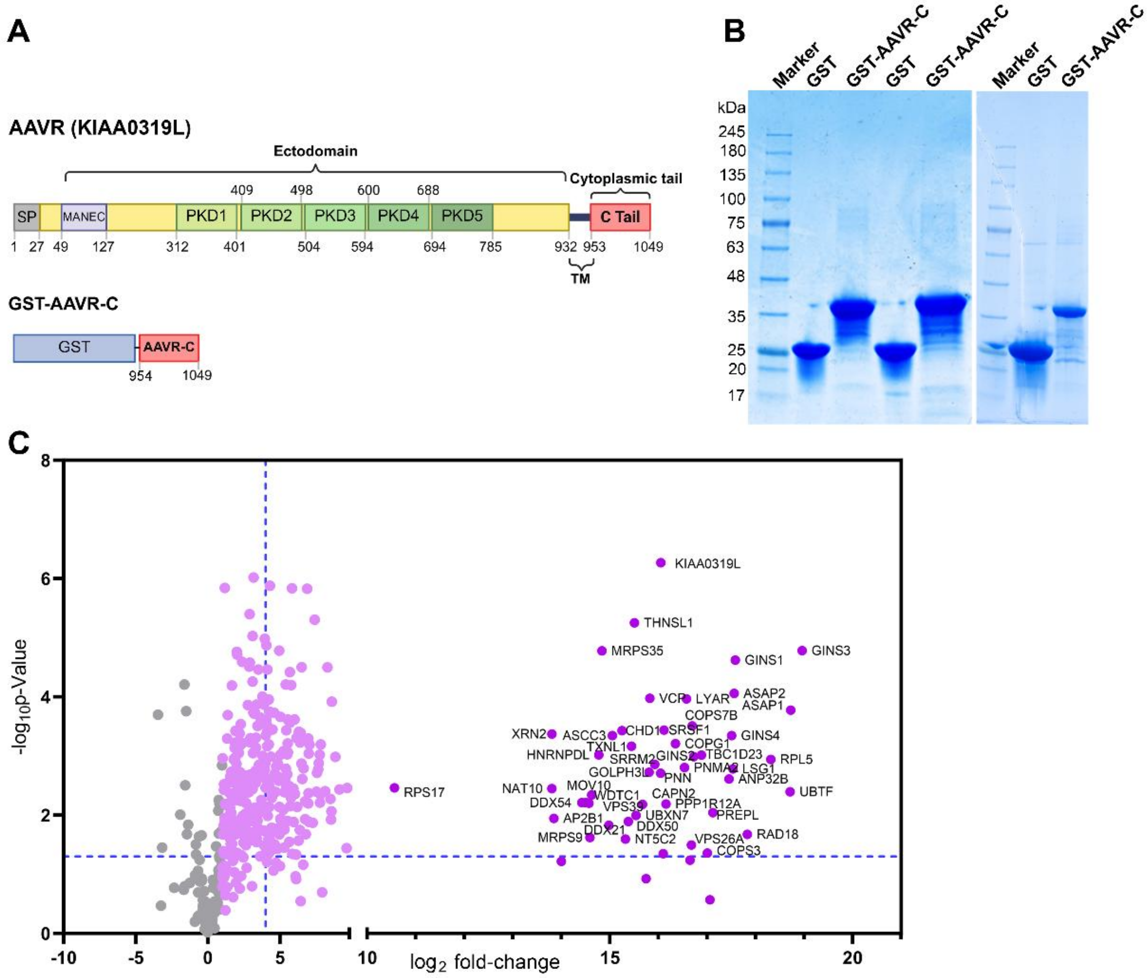


**Figure S2**

**Fig. S2. Identification of TBC1D23 as a cytosolic interactor of KIAA0319L (AAVR).**

**(A) A diagram of AAVR.** Schematic representation of human KIAA0319L (AAVR) domain organization. AAVR is a type I transmembrane protein containing an N-terminal signal peptide (SP), a MANEC domain, five extracellular polycystic kidney disease (PKD) domains (PKD1–PKD5), a single transmembrane helix, and a short C-terminal cytosolic tail. The C-terminal region (AAVR-C; aa954-1049) used for biochemical interaction studies is indicated. **(B) GST pulldown assay.** Coomassie blue–stained SDS–PAGE analysis of recombinant GST or GST-tagged AAVR C-terminal domain (GST–AAVR-C) used for affinity purification. Comparable loading and integrity of purified proteins are shown. **(C) Quantitative mass spectrometry.** Volcano plot of proteins enriched in GST–AAVR-C pulldown relative to GST control, as determined by label-free quantitative LC–MS/MS. Each point represents an individual protein, plotted by $\log_2$ fold enrichment versus $-\log_{10}$ adjusted $P$ value.

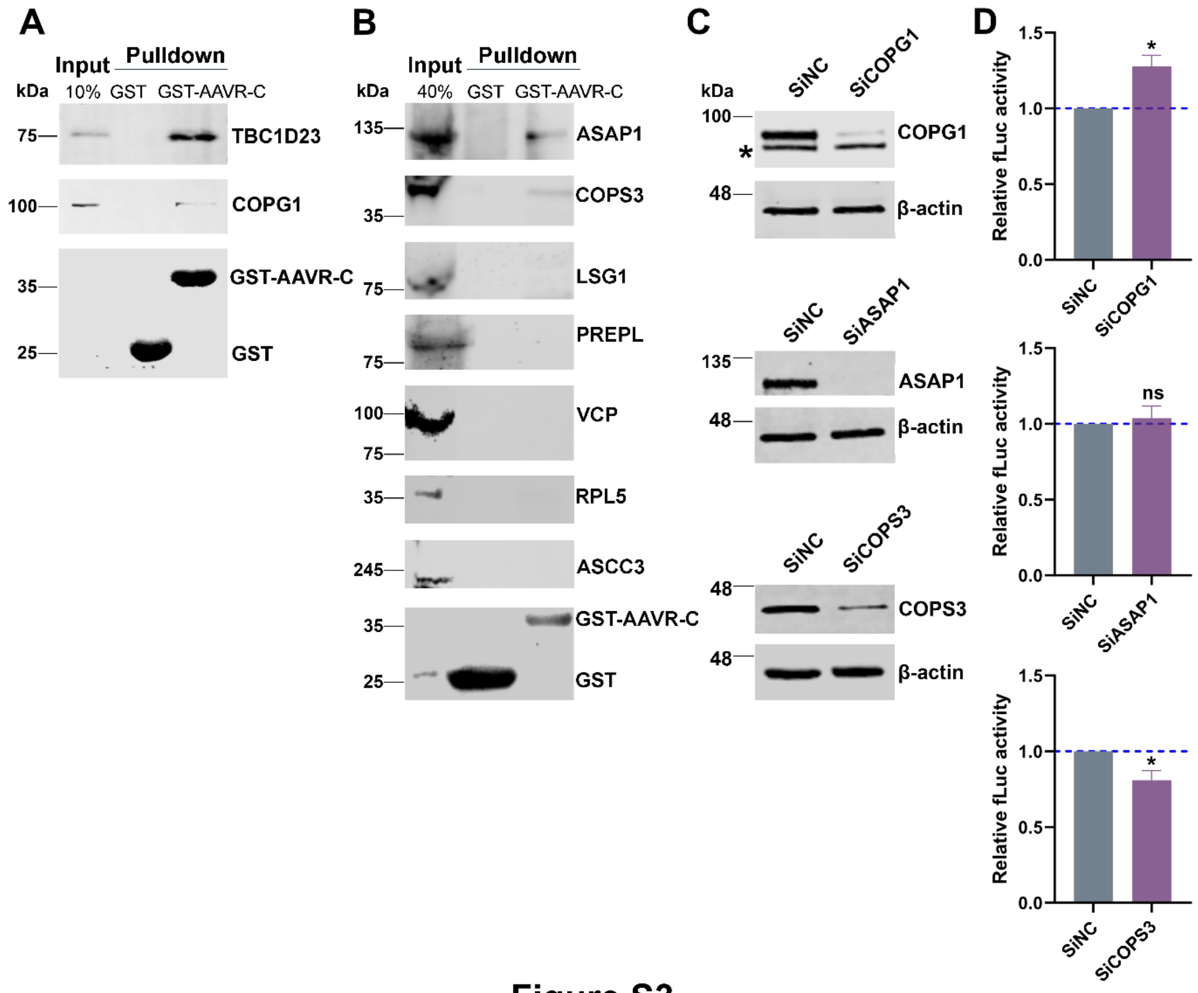


### Figure S3

**Fig. S3. Validation of AAVR-C-interaction with identified candidates and their role in rAAV transduction.**

**(A&B) GST pull-down assay validating selected candidate proteins identified in the AAVR-C interactome screen.** Cell lysates were incubated with GST or GST-tagged AAVR cytoplasmic domain (GST–AAVR-C), and bound proteins were analyzed by immunoblotting using the indicated antibodies. Input lysates and pull-down fractions are shown. **(C) Immunoblot validation of siRNA-mediated knockdown of COPG1, ASAP1, and COPS3 in HEK293 cells.** HEK293 cells were transfected with siNC or the indicated gene-specific siRNA pool. At 48 h post-transfection, cell lysates were analyzed by immunoblotting for the indicated proteins. β-Actin served as a loading control. The asterisk indicates a nonspecific band detected by the COPG1 antibody. Molecular weight markers (kDa) are shown on the left. **(D) Effects of siRNA-mediated knockdown of candidate genes on rAAV transduction.** HEK293 cells transfected with the indicated siRNAs were transduced with rAAV2.5T vectors expressing firefly luciferase (fLuc) at an MOI of 5,000 DRP/cell. Luciferase activity was measured 48 hpt and normalized to the siNC control, which was set to 1. Data represent the mean ± SD of three independent experiments. $P < 0.05$; ns, not significant ($P > 0.05$).

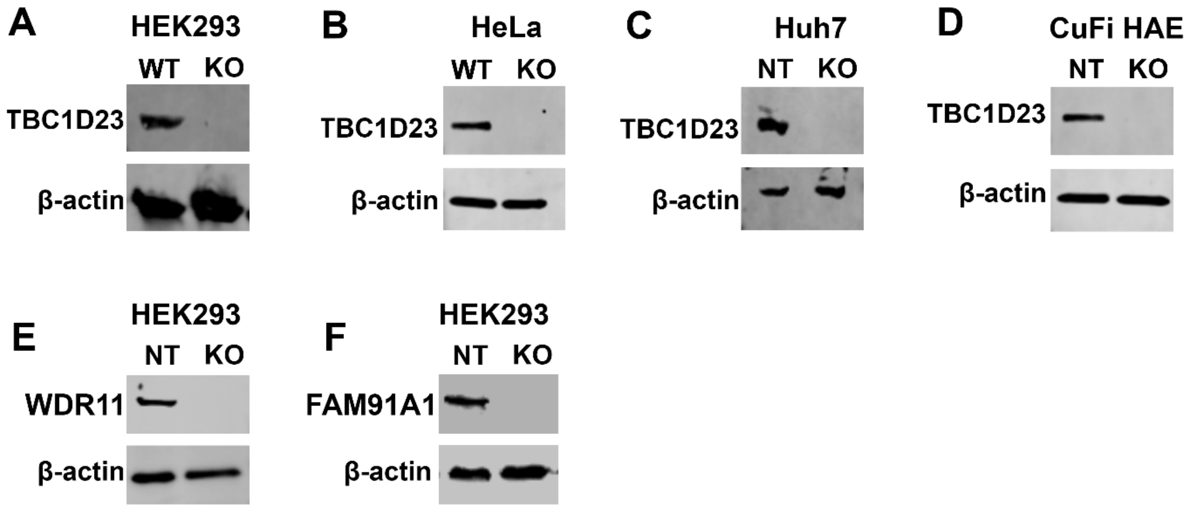


### Figure S4

**Fig. S4. Validation of knockout of TBC1D23, WDR11, and FAM91A1.**

**(A–D) Immunoblot analysis of TBC1D23 expression.** Lysates of wild-type (WT) or non-targeting control (NT) and TBC1D23-KO HEK293 (A), HeLa (B), Huh7 (C), and CuFi HAE (polarized human airway epithelial culture) (D) were analyzed by SDS–PAGE followed by Western blotting using α-TBC1D23. **(E&F) Immunoblot analysis of WDR11 and FAM91A1 expression in NT and WDR11-KO and FAM91A1-KO HEK293 cells.** Cell lysates were analyzed by SDS–PAGE followed by Western blotting using antibodies against WDR11 (E) and FAM91A1 (F). β-actin was used as a loading control.

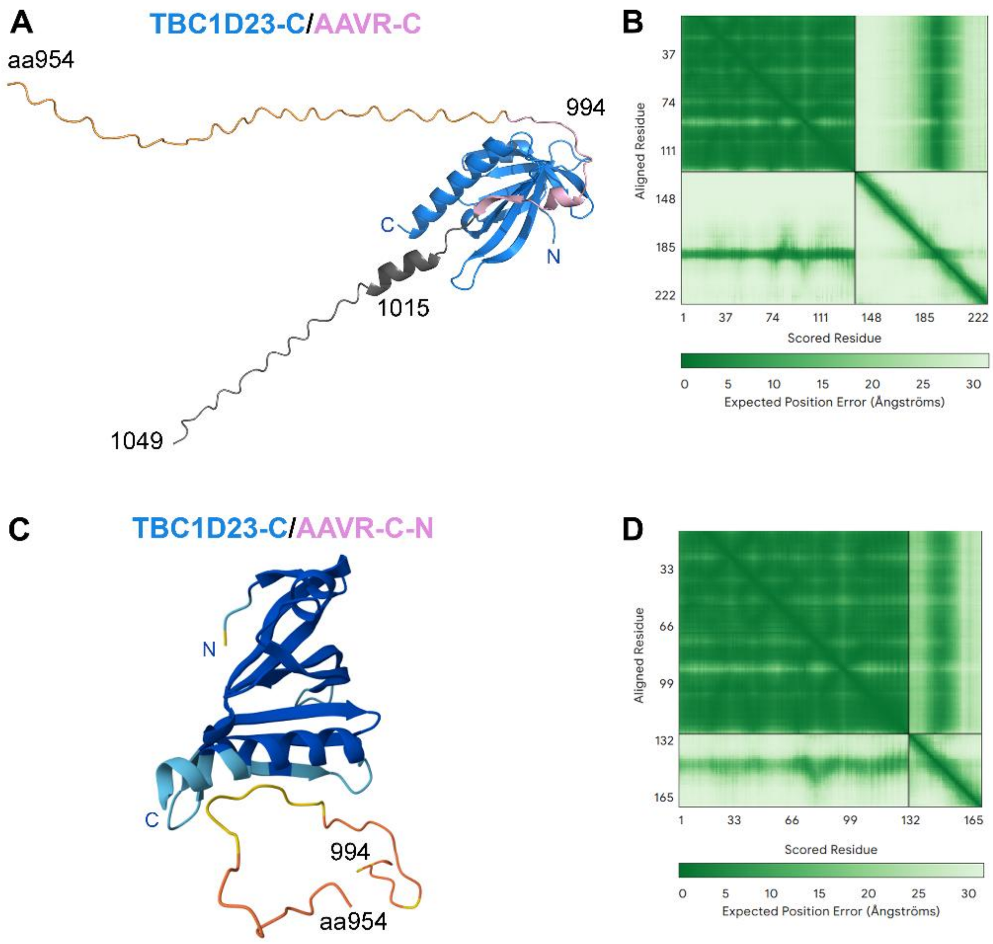


**Figure S5**

**Fig. S5. AlphaFold predictions of interactions between TBC1D23-C and the AAVR C.**

**(A) Predicted complex between TBC1D23 and AAVR-C.** The structure of TBC1D23-C (aa559-684; PDB #6JM5, blue) was used to align with AAVR-C (aa954-1049; magenta) using AlphaFold 3. Acidic residue boundaries aa 994-1015 are indicated. The model shows a structured TBC1D23-C domain with the AAVR-C tail largely disordered. Predicted model confidence scores are shown (ipTM = 0.70, pTM = 0.59). **(B) Predicted aligned error (PAE) plot corresponding to the TBC1D23-C/AAVR-C model in panel A.** Lower PAE values (dark green) indicate higher confidence in relative residue positioning. **(C) Predicted complex between TBC1D23-C (blue) and the N-terminal region of AAVR-C (AAVR-C-N, aa 954-994; brown).** The AAVR segment remains largely unstructured, with limited confidence in intermolecular contacts. Predicted confidence scores are shown (ipTM = 0.38, pTM = 0.71). **(D) PAE plot corresponding to the model shown in panel C.** Lower PAE values (dark green) indicate higher confidence in relative residue positioning.

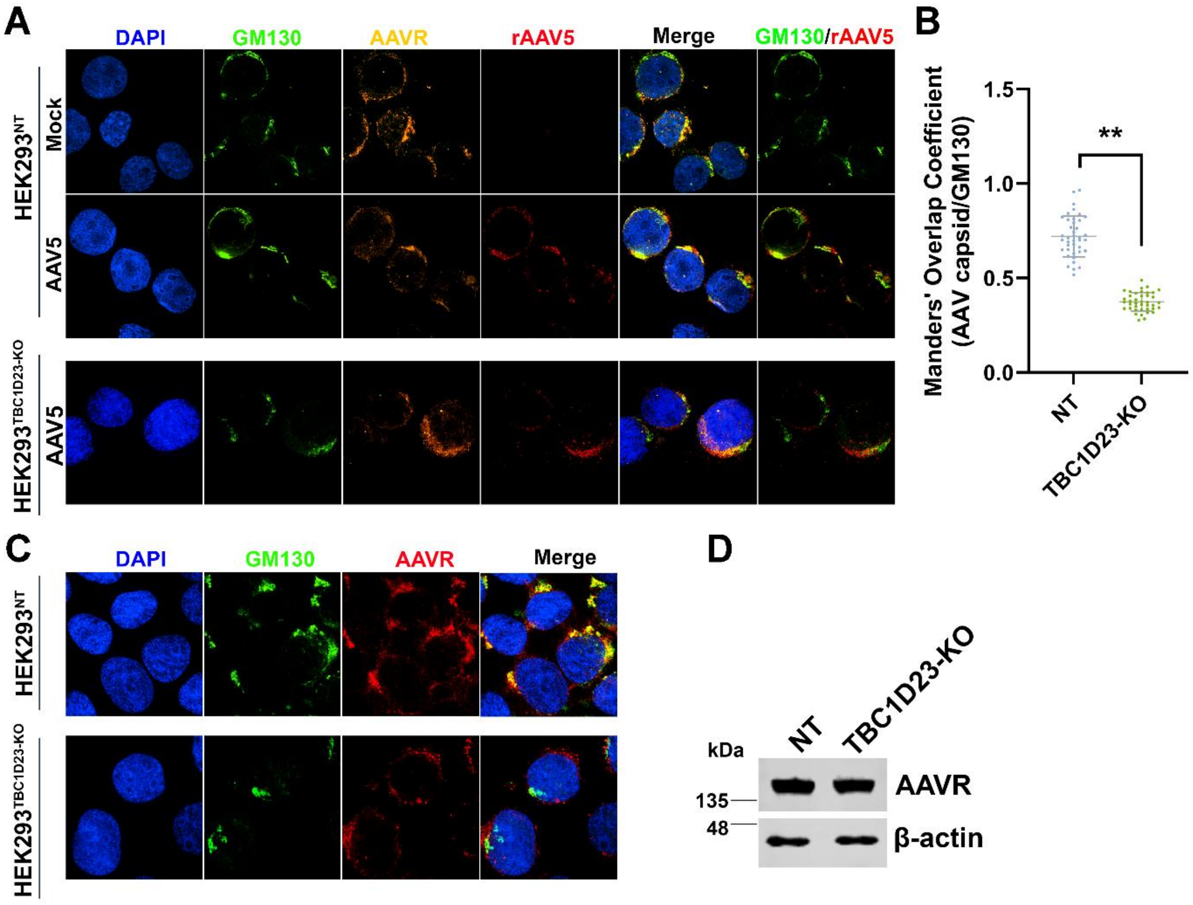


**Figure S6**

**Fig. S6. TBC1D23 knockout alters the Golgi-associated localization of AAV and AAVR without affecting AAVR expression.**

**(A&B) TBC1D23 knockout impairs Golgi localization of the AAV–AAVR complex.** Non-targeting (NT) and TBC1D23-KO HEK293 cells were transduced with rAAV5 at an MOI of 50K. At 6 hpt, cells were immunostained for AAV capsid, the *cis*-Golgi marker GM130, and AAVR, and imaged under a confocal microscope (CSU-W1 SoRa; Nikon) at x 60 magnitude (A). Nuclei were counterstained with DAPI (blue). Merged images are shown on the right. Manders' Overlap Coefficient was calculated for AAV colocalization with the GM103 in single cells (n = 40 per group) using ImageJ (Fiji). Each dot represents one cell; bars show mean ± SD (B). **P < 0.01. **(C) Immunofluorescent staining for endogenous AAVR in TBC1D23-KO HEK293 cells.** HEK293 NT or TBC1D23-KO cells were fixed, permeabilized, and stained for endogenous AAVR (red) and GM130 (green). Nuclei were counterstained with DAPI (blue). Images were acquired using confocal microscopy with a 60× objective. **(D) Immunoblot analysis of endogenous AAVR protein expression in TBC1D23-KO cells.** NT and TBC1D23-KO cells were analyzed by Western blotting for AAVR expression. β-actin served as a loading control.

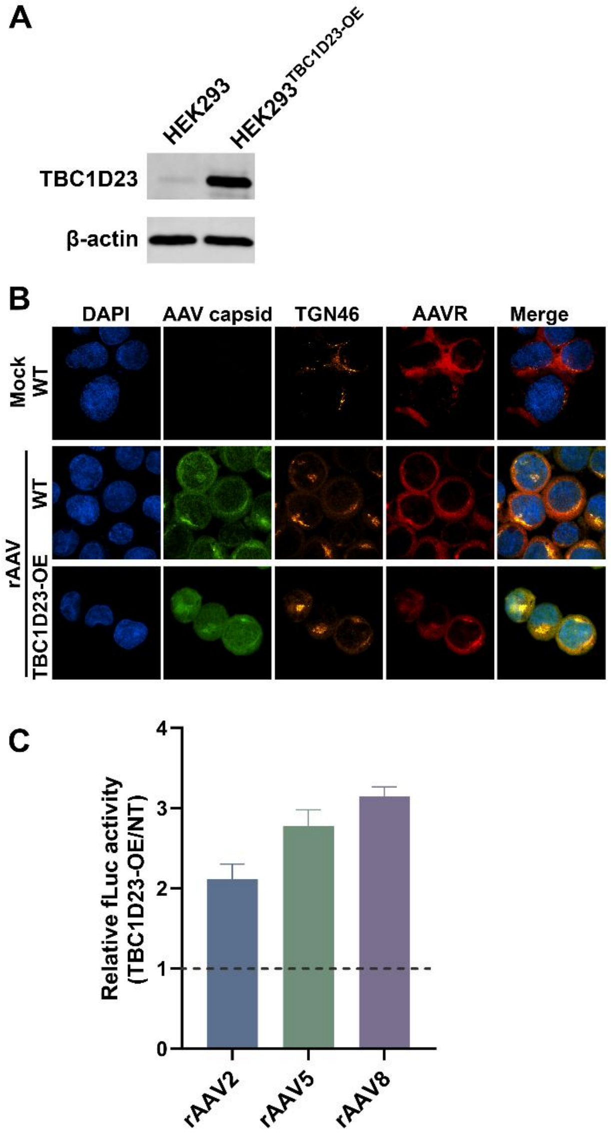


**Figure S7**

**Fig. S7. Overexpression of TBC1D23 increases nuclear localization of AAV and transduction of rAAV.**

**(A) Overexpression (OE) of TBC1D23 in HEK293 cells.** Whole-cell lysates from parental HEK293 cells and the $HEK293^{TBC1D23\text{-}OE}$ cells stably overexpressing TBC1D23 were analyzed by SDS–PAGE followed by Western blotting using α-TBC1D23. β-actin served as a loading control. **(B) Immunofluorescence analysis of AAV capsid trafficking in HEK293 cells**. WT and $HEK293^{TBC1D23\text{-}OE}$ cells were mock-treated or transduced with rAAV5 at an MOI of 20K. At 6 hpt, cells were stained for nuclei (DAPI, blue), AAV capsid (green), the *trans*-Golgi network marker TGN46 (orange), and AAVR. Merged images are shown on the right. Representative images illustrate comparable intracellular localization patterns across conditions, but an increase in nuclear-imported capsids. **(C) Transgene expression.** WT and $HEK293^{TBC1D23\text{-}OE}$ cells were mock-treated or transduced with rAAV2, rAAV5, or rAAV8 at an MOI of 20K. At 48 hpt, fLuc activity was quantified. Relative fLuc activity normalized to WT is shown with mean ± SD (n = 3 independent experiments).

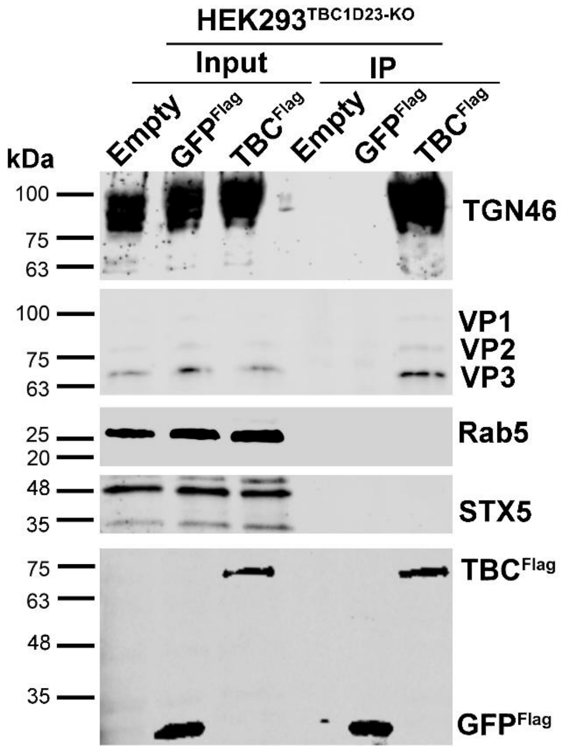


#### Figure S8

**Fig. S8. Endosomal immunoprecipitation (Endo-IP) assays using TBC$^{Flag}$- and control GFP$^{Flag}$-expressing HEK293$^{TBC1D23\text{-}KO}$ cells.**

HEK293$^{TBC1D23\text{-}KO}$ cells were reconstituted with WT TBC1D23$^{Flag}$ (TBC$^{Flag}$) or GFP$^{Flag}$ (control) were transduced with rAAV5 at an MOI of 50K. At 2 hpt, cells were washed, lysed under detergent-free conditions by Dounce homogenization, clarified, and Flag-tagged vesicles were immunoprecipitated using anti-Flag M2 magnetic beads. Inputs (10%) and IP eluates were analyzed by immunoblotting for the indicated markers (TGN46, STX5, and Rab5), AAV capsid proteins (VP1/VP2/VP3), and TBC1D23$^{Flag}$ or GFP$^{Flag}$. Input: 10% of the cell lysates used for IP. Protein ladders are indicated to the left. Input represents 10% of the total cell lysate used for immunoprecipitation. Protein size markers are indicated on the left.

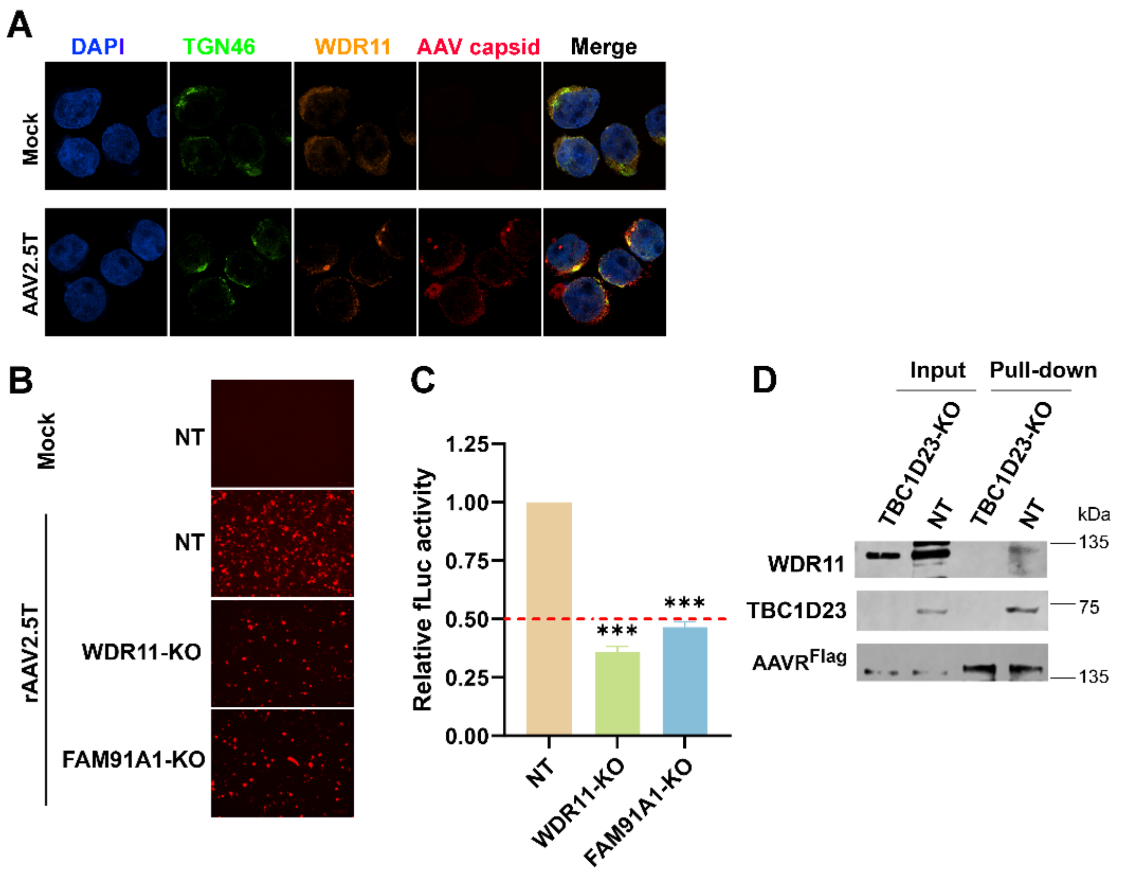


#### Figure S9

**Fig. S9. The WDR11-FAM91A complex is required for efficient rAAV transduction. (A) WDR11 is associated with the AAV capsid at the TGN.**

HEK293 cells were transduced with rAAV2.5T at an MOI of 10K. At 4 hpt, the cells were stained for TGN46, WDR11, and AAV capsid. Images were taken under a confocal microscope at a magnification of x 60. **(B&C) rAAV2.5T transduction of WDR11-KO and FAM91A-KO HEK293 cells.** Non-targeting (NT), WDR11-KO and FAM91A-KO HEK293 cells were transduced with rAAV2.5T at an MOI of 10K. At 2 dpt, cells were imaged for mCherry expression (B) and assayed for fLuc (C). (B) Representative fluorescence and brightfield images of HEK293 cells are shown. (C) Quantification of fLuc activity. Data is normalized to NT control. Statistical significance was assessed by one-way ANOVA with multiple comparisons (***$P < 0.001$; ****$P < 0.0001$). **(D) WDR11 association with the AAVR complex is dependent on TBC1D23.** Co-immunoprecipitation (Co-IP) examines the association of WDR11 with the AAVR complex in AAVR$^{Flag}$ overexpressing NT and TBC1D23-KO HEK293 cells. Cell lysates were immunoprecipitated using anti-Flag antibody, and the precipitated fractions were analyzed by SDS–PAGE followed by immunoblotting for WDR11 and TBC1D23. Comparable levels of WDR11 were present in input samples (10%), whereas WDR11 recovery in the precipitated fraction was absent in TBC1D23-KO cells, consistent with a role for TBC1D23 in facilitating WDR11 association with AAVR.
